# Impact of multiple phosphorylations on the tau-R2/tubulin interface

**DOI:** 10.1101/2025.02.19.638976

**Authors:** Jules Marien, Chantal Prévost, Sophie Sacquin-Mora

## Abstract

The phosphorylation of the microtubule-associated tau protein plays a key role in the regulation of its physiological function. In particular, tau hyperphosphorylation affects its binding on the tubulin surface, destabilizing the tau-microtubule interface and leading to the accumulation of fibrillar aggregates in the brain. In this work, we performed classical Molecular Dynamics simulations for the tau-R2/tubulin assembly with various phosphorylations states of serines 285, 289 and 293. We analyze the resulting trajectories to obtain a detailed view of the protein interface in the complex, and the impact of tau phosphorylations on the stability of this assembly, and on the tubulin disordered C-Terminal tails (CTTs) mobility. We show how the tubulin CTTs help maintaining the tau-R2 fragment on the tubulin surface despite the destabilizing effect induced by phosphorylations. Conversely, tau phosphorylation affects the CTTs flexibility and their potential activity as MAP recruiting hooks. Furthermore, counterion mediated bridges between the phosphate groups and tubulin glutamates also contribute to the binding of tau-R2 on the MT. Overall, the complex dynamics of this fuzzy phosphorylated assembly sheds a new light on the importance of the cytoplasmic environment in neurons in the context of Alzheimer’s disease.

## 1. Introduction

Microtubules (MTs) are a key component of the eukaryotic cytoskeleton,^1^ whose assembly and activity is regulated by a number of microtubule-associated proteins (MAPs). In neurons, MTs interact with a broad family of « classical » MAPs, such as MAP-2, MAP-4 and tau. The later is the most abundant, as it constitutes more than 80% of the neuronal MAPs.^2, 3^ Tau is a long (over 400 amino acids) intrinsically disordered protein, which controls MT assembly spatial organization and stability,^4-6^ and the motility of motor proteins along MTs.^7-10^ The tau/MT binding pattern is complex and dynamic, as it involves the disordered, highly flexible, C-terminal tails (CTTs) of tubulin.^11, 12^ Tubulin CTTs are glutamate rich, highly charged, intrinsically disordered regions (IDRs), and therefore do not appear in tubulin structures produced by cristallography or cryo-EM. α- and β-tubulins exist as multiple variants (named isotypes) that differ mainly by their CTT length and sequence. The CTTs also have a functional role,^13^ and act as fine regulators of MT function,^14-16^ modulating MT assembly,^17^ dynamics^18-20^ and interaction with motor proteins^21, 22^ and MAPs.^23^ The tau/ MT assembly thus forms a dynamic, fuzzy complex,^24^ involving both ordered and disordered elements, which remains difficult to characterize with structural biology approaches.

Furthermore, Tau is a target for numerous post-translational modifications (PTMs).^25-27^ In the healthy brain, tau reversible phosphorylation plays a role in MT regulation.^28, 29^ Tau hyperphosphorylation however, has been connected to several neurodegenerative diseases (tauopathies), such as Alzheimer’s disease (AD), that involve fibrillar aggregation.^30, 31^ Over the 85 potential phosphorylation sites of tau (which mostly include serine and threonine residues), about 40 of these were only observed in AD brains.^32^ The question remains to determine how these PTMs will impact the interaction between tau and the MT surface.^33-35^

Using a combination of Cryo-EM data and modeling approaches on the tau-R2/tubulin assembli,^36^ Brotzakis et al. identified a stongly interacting region nest to the C-terminal part of the tau-R2 fragment, while its N-terminal part was weakly interacting with the tubulin surface. In an earlier work,^37^ we characterized the interface between the tau R2 domain (which belongs to the protein central MT binding domain) and tubulin, using homology modeling and all-atom Molecular Dynamics (MD) simulations. In particular, our study highlighted the key role played by the three serine residues of tau-R2 (Ser285, Ser289, and Ser293), which lie in the center of the R2 strongly interacting region defined by Brotzakis et al.^36^ In addition, we investigated the complex interplay between tau-R2 and tubulin CTTs. On one hand, we showed how these CTTs contribute to stabilizing the tau/tubulin interface. On the other hand, we also showed how tau binding on the MT surface impacts the CTTs mobility, and might therefore contribute to regulating their activity as hooks that are responsible for the recruitments of MAPs.

The present study investigates the impact of multiple phosphorylations of the R2 serine residues on the tau/tubulin interface for a cleaved tubulin trimeric system without CTTs, and for the βI/αI/βI and βIII/αI/βIII tubulin isotypes. As could be expected, tau phosphorylation leads to a destabilization of the interface. However, the interaction of phosphorylated tau-R2 with the charged residues from the tubulin CTTs, notably via sodium mediated counter-ion bridges, can contribute to weaken this destabilizing effect. Furthermore, we use new metrics specifically tailored to describe IDPs and IDRs^38^ to characterize how tau -R2 multiple phosphorylations impact the mobility and flexibility of the tubulin CTTs.

## 2. Material and Methods

### Starting models and phosphorylation states

All starting models were derived from our earlier work on the R2-tubulin assembly.^37^ Briefly, the PDB structure 6CVN^39^ was selected to produce trimers of βI/αI/βI and βIII/ αI/βIII tubulin isotypes (see Tables S1 and S2 for the selected tubulin sequences) in complex with the R2 repeat domain of the Tau protein using Modeler v10.2.^40, 41^ For each isotype combination, 100 models were generated differing by their tubulin CTTs conformations, and three models were selected as starting points to launch independent replicas. An additional system with cleaved CTTs was designed by removing the CTTs from the βI/αI/βI starting structures. Note that the terminal residues of tau-R2 were kept charged, which might influence electrostatic interactions with the tubulin surface compared to full-length Tau. The residues of the entire complex were assigned their canonical protonation state at pH 7. No disulfide bond was enabled, GTP/GDP and Mg were excluded from the complex, with no noticeable destabilization (see Figure S4 in ref. 37). More details regarding the simulation setup are available in the *Material and Methods* section of our previous work.^37^

We term *P-state* the specific combination of phosphorylated residues in the complex. Since there are three serines in the R2 domain (Ser285, Ser289 and Ser293), there are eight possible P-states (termed P000, P001, P010, P100, P011, P101, P110 and P111, with 0 indicating a canonical serine and 1 a phosphoserine) for the tau fragment (see Figure 1 for a snapshot of the P111-state of the βI/αI/βI isotype). The CHARMM-GUI web-server (https://www.charmm-gui.org/)^42^ was used to generate all possible P-states for all the earlier mentioned starting models. The SP2 patch was used to obtain the dianionic form of phosphoserines, which is most common in solution at pH 7.^43^ Table S3 provides a summary of all the systems built for the present study and the total number of simulations is therefore 72 (two isotypes combinations and a tubulin trimer without CTTs, eight P-states, and three replicas for each system/P-state combination).

**Figure 1.**
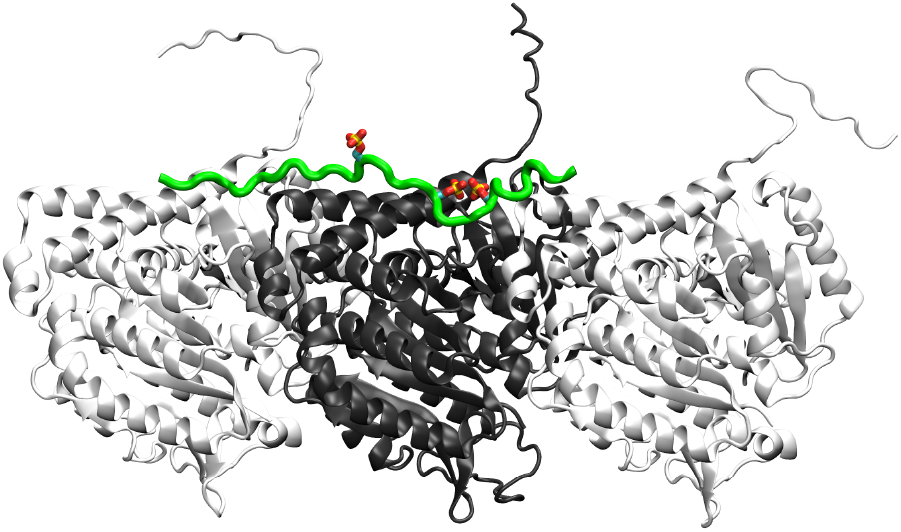
Starting structure for the tau-R2/tubulin assembly of isotype βI/αI/βI in the P111 state. The tubulin monomers are shown in dark and light grey for the α and β subunits respectively, while tau-R2 is shown in green. Phosphorylations on Ser285, 289 and 293 are displayed in stick representation. Figures 1, 7, 8 and 9 were prepared using Visual Molecular Dynamics.^56^

### All-atom molecular dynamics simulations

Solvation in TIP3P water^44^ was obtained with the CHARMM-GUI server neutralizing the rectangular box and adding a concentration of 0.15 mol.L^-1^ of NaCl. Histidines were unprotonated in the HID state, and the protonation of other amino acids was set to pH 7. All-atom molecular dynamics simulations were carried out with NAMD v2.14^45^ and the CHARMM36m^46^ force field. The SHAKE algorithm was used to fix hydrogen covalent bonds,^47^ and periodic boundary conditions were enabled. The Nose-Hoover Langevin piston algorithm set the pressure to 1 atm, and Langevin dynamics maintained the temperature to 300 K.^48, 49^ Electrostatic interactions were calculated using the Particle Mesh Ewald method^50^ with a grid space of 1 Å and an interpolation order of 6. The nonbonded cutoff was 1.2 nm. The integration time step was 2 fs and frames were saved every 100 ps in production. To make sure that the system would not collide with its periodic image along the smaller side of the box, we constrained its overall orientation with a harmonic restraint on the α-carbons of the tubulin cores. The trimer is otherwise free to deform. The scaled normalized force constant was set to 0.05 kcal.mol^-1^ and the restraint was implemented with the ColVar module available in NAMD.^51, 52^

The equilibration procedure consisted of 10,000 steps of energy minimization followed by 2 ns of equilibration with constrained protein heavy atoms in the NVT ensemble. Production lasted for 200 ns for each replica, resulting in 600 ns of cumulative simulated time for each system.

### Analysis of the MD trajectories

All systems were aligned in post-processing on the tubulin cores. Ions were then wrapped around the R2 repeat domain to allow for nP-collab^38^ analysis. Ad-hoc python scripts using the MDTraj^53^ and MDAnalysis^54^ modules were used to perform analyses on the complete (200 ns for each replica) MD trajectories. The fraction of native contacts Fnat is defined as the fraction of conserved contacts between two selections at time t with respect to the contacts between the two selections in the first frame. In this study we usually define one selection as the R2 peptide and the other selection as the tubulin monomers, unless specified otherwise. MM/PBSA calculations of the binding enthalpies between tau R2 and the tubulin heterotrimer were ran on the last 150 ns of each production run with a stride of 10 frames using a combination of the APBS program^55^ and the Cafe_plugin on VMD.^56, 57^ Proteic Menger Curvatures (PMCs), Local Curvatures (LCs) and Local Flexibilities (LFs) were defined and calculated as per our previous work on nP-collabs using the following script: https://github.com/Jules-Marien/Articles/tree/main/_2024_Marien_nPcollabs/Demo_and_scripts_Proteic_Menger_Curvature.^38^

## 3. Results and discussion

### Conformational variability of the tau-R2/tubulin interface upon phosphorylation

We used the R2 non-phosphorylated state as a reference for the comparison with the different P-states. Following the work of Brotzakis et al,^36^ we define the R2Val287-R2K294 and R2Lys274-R2Lys281 fragments as the strong and weak tubulin binding sites of tau-R2, respectively. Timewise RMSD of the R2 peptide were calculated using the first frame as reference (see Figure S1). As expected, the RMSD is very high (above 5 Å) for all systems, attesting for a highly-dynamic disordered behavior.

We first consider the changes in the contact maps for the tau-R2/tubulin interface upon adding phosphorylations to R2, our reference states corresponding to Fig. 4 in Ref. 37. For the triply phosphorylated systems, the contact variation maps shown in Figure 2 reveal an overall loss of contacts (in blue) for both the weak and strong binding sites. Most contact losses are centered around R2-Gln288 and R2-Ser289, the latter forming the core of the strong tubulin binding site.^37^ Strikingly, contact time can weaken by nearly 80% in the strong binding site, suggesting that phosphorylations greatly perturb the interface. In addition, the contact loss between R2 and the αI-CTT for the βI/αI/βI tubulin isotype (Figure 2b) is similar to what was observed by Dasari et al. when investigating the phosphorylation of the tau-R4 fragment bound to tubulin.^58^

**Figure 2.**
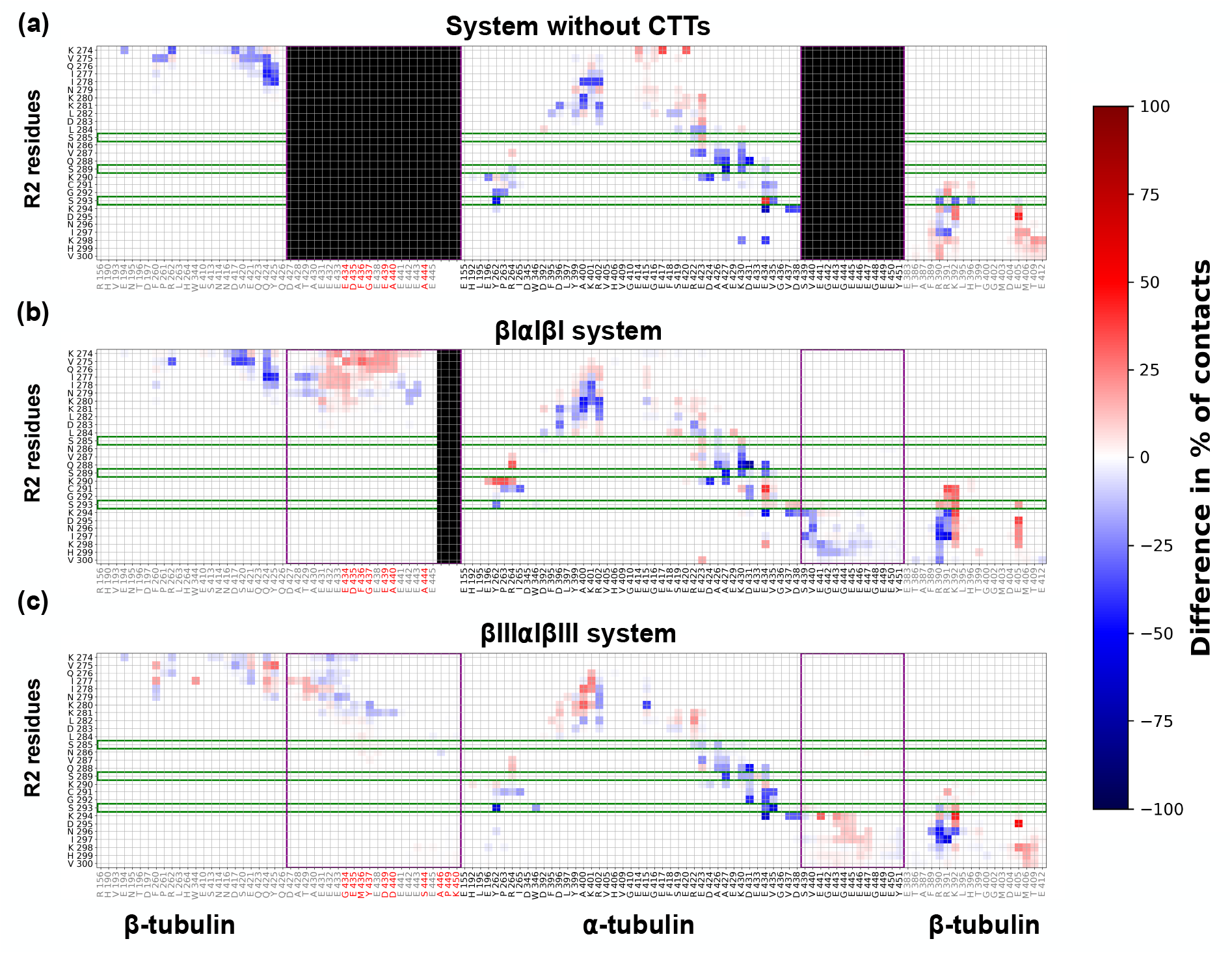
Variations in the tau-R2/tubulin contact maps of the P111 systems compared to the P000 systems. For each system, the datat of the three replicas were concatenanted. The tubulin CTTs for the βI/αI/βI and βIII/αI/βIII isotypes (central and lower panels) are delimited by a purple frame, while the phosphorylated serines (Ser 285, 289 and 293) of tau-R2 are highlighted by green frames. On the horizontal axis, α- and β-tubulin residues are displayed in black and grey respectively. (a) Tubulin without CTTs (b) βI/αI/βI trimer (c) βIII/αI/βIII trimer. Blue areas highlight a contact loss upon phosphorylation, while the red areas highlight a contact increase.

**Figure 3.**
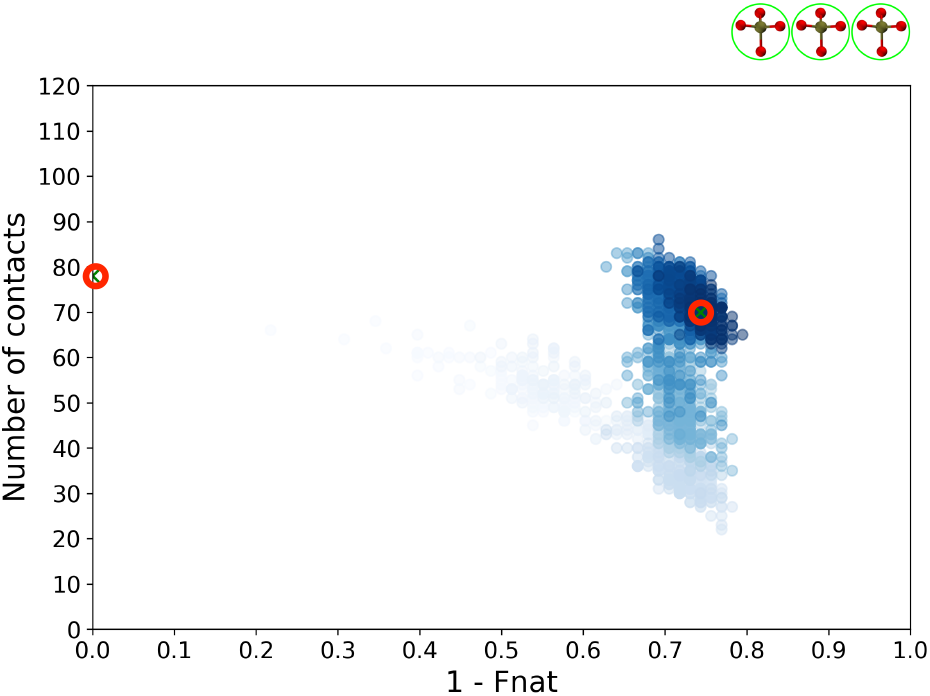
Number of R2/tubulin contacts for the third replica of the βI/αI/βI trimer in the P111 state, as a function of the native contacts loss (1-Fnat). The red circles highlight the initial and final points of the MD trajectory, and the blue dots grow darker along time.

**Figure 4.**
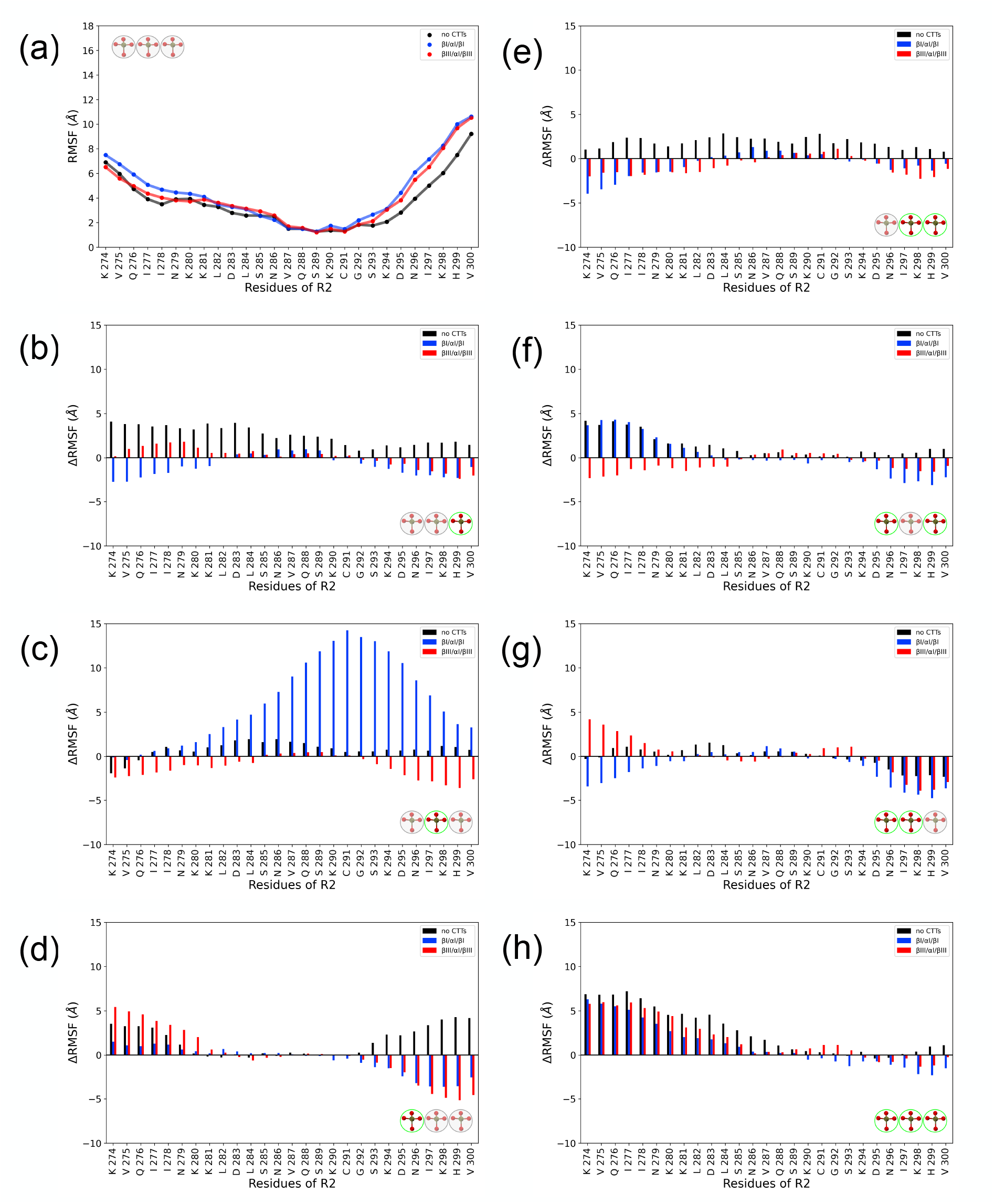
(a) RMSF of tau-R2, from the concatenation of the three simulations for each system in the unphosphosphorylated state. (b-h) RMSF variation upon phosphorylation, the scheme in the lower right corner highlights the considered P-state, from P001 (panel b) to P111 (panel h).

Considering all P-states (available in Figure S2), a general trend can be identified. While the strong and weak binding sites do display smaller contact times with the tubulin surface, new contacts are also formed between the tubulin CTTs and R2, limiting the overall contact loss. This is visible in Figure 2b for the first βI-CTT of the βI/αI/βI tubulin trimer, and in Figure 2c, for the αI-CTT of the βI/αI/βI tubulin trimer. In Figure 3, which displays the total number of R2/tubulin contacts for the third replica of the βI/αI/βI isotype in the P111 state as a function of the native contacts loss (1-Fnat), the blue dots grow darker along time. After an initial phase where both contact number and Fnat steadily decrease, the Fnat value stabilizes in the 20-25% range, while the total contact number increases to reach a final value that is very close to the starting one. This highlights the formation of new contacts along time, and the fuzzy nature of the complex. A similar behavior is observed across all systems and most replicas, as can be seen on Figure S3.

We then calculated the RMSF on the Cαs between the starting conformation of R2 for each type of system and its conformations along the corresponding replicas. We already characterized that the strong binding site is stable along all non-phosphorylated trajectories^37^ (see Figure 4a), and that the N-ter and C-ter parts of R2 present similar behaviours for the tubulin without CTTs and the βI/αI/βI and βIII/αI/βIII tubulin isotypes. We calculated the RMSF variation between the unphosphorylated and all the phosphorylated states (see Figure 4b-h), which allows us to probe the evolution of the interface. The triple pSer285/pSer289/pSer293 phosphorylation does not disturb the strong binding site as the ΔRMSF for the P111 state remains below 2Å (see Figure 4h). We even notice a slight stabilization of the C-terminal part of R2 when the αI-CTT of tubulin is present. On the contrary, the weak binding site is more mobile for every system, with twice as much difference for the system with cleaved CTTs. These observations are valid for most P-states, although phosphorylations of Ser289 and/or Ser293 tend to increase the ΔRMSF of the strong binding site more than phosphorylation of Ser285. A noticeable event can be highlighted for the P010 state, where the ΔRMSF skyrockets over 1 nm around the strong binding site (see Figure 4c and Figure S1b). We indeed observed a complete separation of R2 from its original interface for one of the replicas. The complex is however in part rescued by the βI-CTT on the N-terminal side of R2, which keeps the latter from fully dissociating. These elements underline the possible roles of tubulin CTTs as complex stabilizers and recruiters.

### Energetic considerations

We computed the binding enthalpies between tau-R2 and the tubulin trimer with the MM/PBSA method in order to understand the energetic contribution of the addition of phosphorylations (see Figure 5, and Table S4 and Figure S4 for the detail of the different contributions). We insist on the limits of such calculations on fuzzy complexes: as is immediately noticeable, the standard deviation on every quantity is significant, not only due to a possible undersampling, but mostly because the complex displays large conformational diversity. One should only consider mean values relative to one another and not in an absolute manner. Overall the R2/tubulin binding enthalpy tends to increase with the number of phosphorylations in R2, thus highlighting the weakening effect of this PTM on the interface, in particular when Ser289 is phosphorylated. Another destabilizing factor is the absence of CTTs, as the cleaved systems always present a larger binding enthalpy compared to the uncleaved ones, with the exception of P-states P011 and P111 (see Figure 5). This concurs with our earlier work on the P000 state,^37^ and previous experimental studies,^24, 59^ which showed the stabilizing effect of the tubulin CTTs on the tau/tubulin assembly.

**Figure 5.**
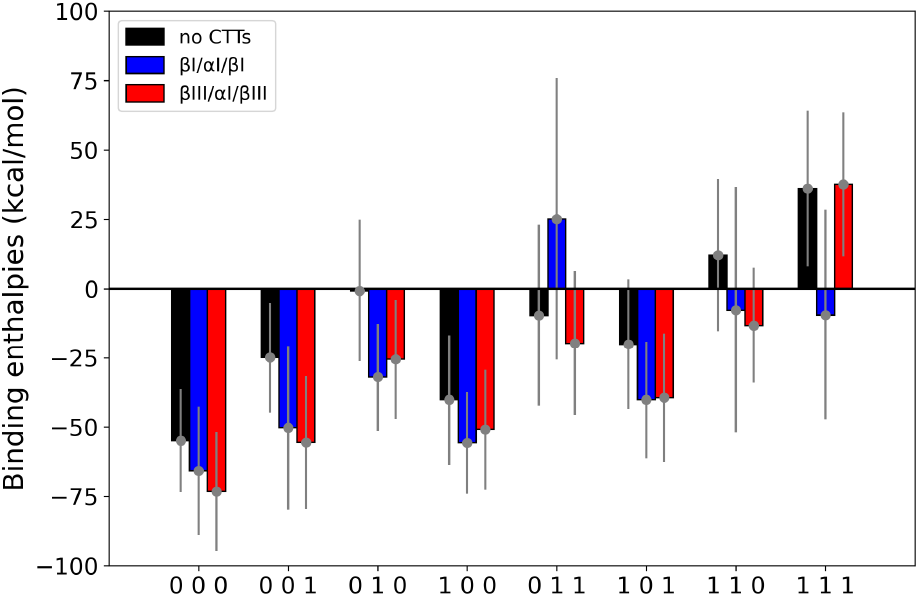
Binding enthalpies for all the P-states in the three systems between the R2 repeat domain and the tubulin monomers. Calculations were performed with a stride of 10 frames on the last 150 ns of trajectory, and the results from the three replicas were combined for each system. The standard deviation represents the deviation on all calculated values.

In four cases (P011 for βI/αI/βI, P110 for cleaved CTTs, and P111 for cleaved CTTs and βIII/αI/βIII), the binding enthalpy even becomes positive, above 10 kcal.mol^-1^. Unless compensated by a strong change in the entropic contribution, this should signal the complex dissociation. The entropic term is notoriously difficult to estimate for fuzzy complexes as disordered proteins tend to not respond well to standard entropic estimators,^60, 61^ and therefore we did not attempt its calculation. However, since the ΔRMSF profiles for the βI/αI/βI isotype in the P001 and P011 states are very similar (see Figure 4b and e), we can assume that their entropic terms are not so different either, while their enthalpic profiles differ by more than 50 kcal.mol^-1^. In addition, the positive binding enthalpies obtained for the double and triple P-states in Figure 5 do not concur with the low ΔRMSF values shown for the strong binding site in Figures 4e-h. Note that similar results (important increase of the binding free energy and low ΔRMSF) were obtained by Man et al.,^62^ even though they used a different force-field (ff14SB), and their model of the tubulin-R2 assembly did not include the tubulin CTTs. Thus we concluded that another phenomenon must be at play to explain such a striking discrepancy between these two observations.

### nP-collabs contribute to stabilize the interface

In a previous study, we characterized – within the context of the CHARMM36m forcefield - the capacity of sodium ions to bridge phosphate groups.^38^ We called this phenomenon a *n*P-collab, where *n* is the number of phosphates bridged by a single cation. We hypothesized that *n*P-collabs might explain the aforementioned discrepancy and monitored the presence of sodium ions close to each serine, phosphorylated or not (see Figure 6 and Figure S5 for the complete set of trajectories). In the absence of phosphorylations, the proximity of sodium is sporadic at best, with fewer than 50 isolated events over the course of a full 200 ns trajectory for all sodium and all serines (Figure 6a). A single phosphorylation is enough to draw a sodium ion to its vicinity for prolonged periods of time (more than 20 ns), as can be seen in Figure 6b (see also Figure 7a for a snapshot). Addition of two phosphorylations is not sufficient to systematically induce the formation of 2P-collabs since the βIII/αI/βIII system with pSer285 and pSer289 does not display any. However, all systems with pSer289 and pSer293 were able to capture at least two sodium ions in a 2P-collab for 100 ns or more (cf. the dark blue lines in Figure 6c, left panel, and Figure 7a). All simulations of the P111 state exhibit various degrees of 2P-collabs, even briefly reaching 3P-collabs for one replica in the βIII/αI/βIII system (red dots in Figure 6d, central panel, and Figure7b). Closer inspection of the trajectories also allows us to identify the formation of a “stabilizing tripod” mediated by a sodium ion, involving R2-pSer289, R2-pSer293 and αI-Glu434. The sodium ion appears to be stabilizing the conformation of R2 by bridging the phosphoserines in a 2P-collab, and allows for αI-Glu434 to remain a stabilizing residue by mediating the interaction with the phosphoserines (see Figure 7c). This behavior reconciles the low ΔRMSF with the contact losses and the highly positive binding enthalpies. Indeed, since the contacts are now in part mediated by sodium cations, direct contacts do drop compared to the non-phosphorylated interface. MM/ PBSA calculations do not include the solvent -and therefore the cations-explicitly as it is accounted for by the Poisson-Boltzman term. As a result, the proximity of the doubly-charged phosphate groups to the negatively charged surface of the tubulin results in high binding enthalpies since the MM/PBSA method cannot include the nP-collab phenomenon. We hypothesize that a similar phenomenon took place in the earlier mentioned study of Man et al.,^62^ as their binding free energy calculations do not take into account the counter ions in the system.

**Figure 6.**
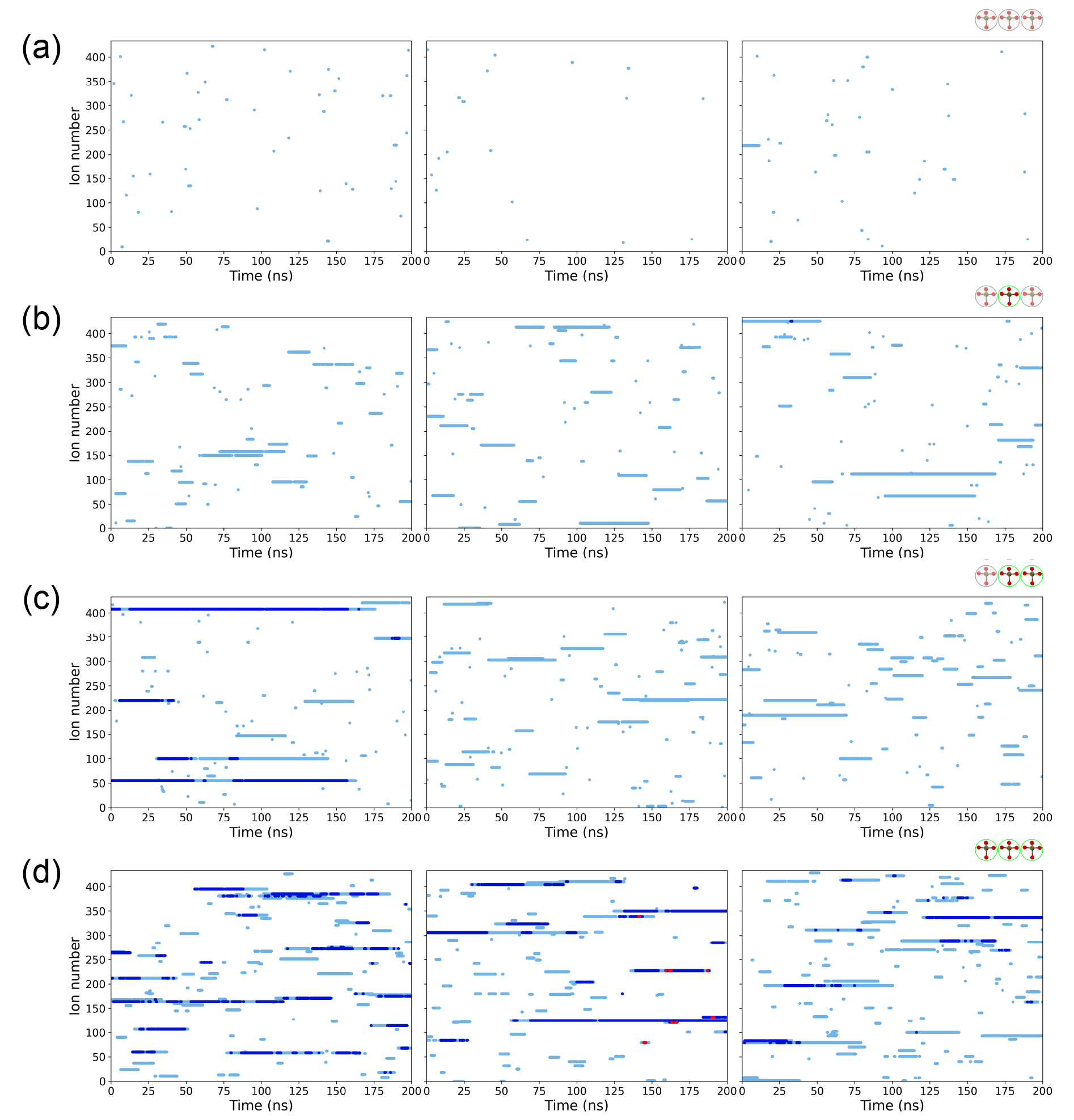
Monitoring the presence of Na^+^ counterions in the vicinity of the R2 serines along the MD trajectories for each replica of the βIII/αI/βIII system. Each horizontal line corresponds to one individual cation. Light blue dots indicate that the cation is less than 4 Å away from a P atom, dark blue indicate proximity to two P atoms (2P-collabs) and red proximity to the three P atoms (3P-collabs).

**Figure 7.**
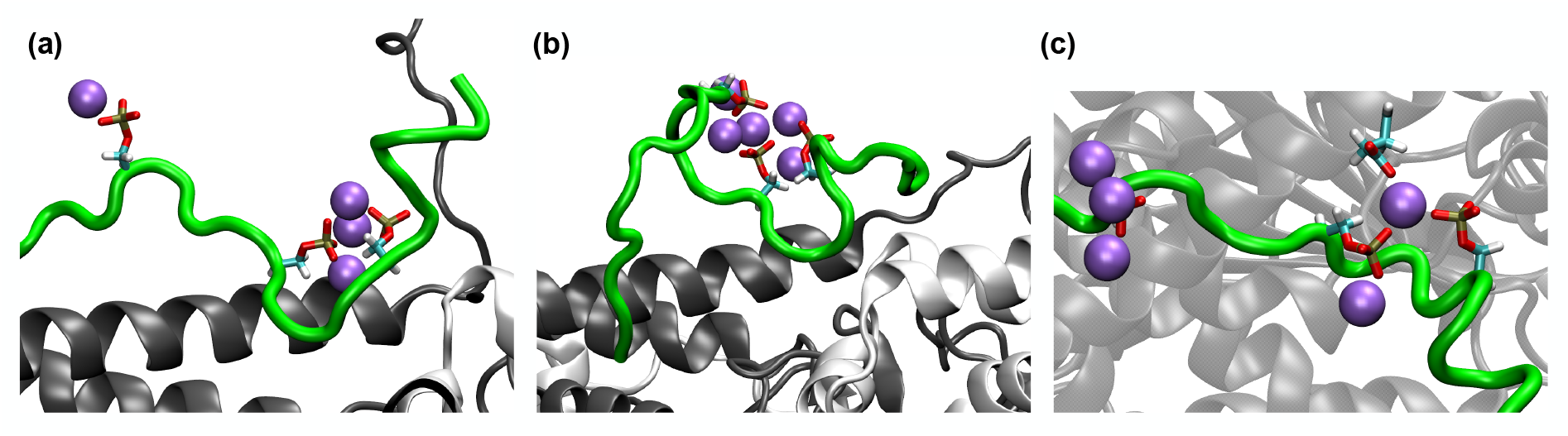
Snapshots from the MD trajectories of the βIII/αI/βIII system in the P111 state highlighting the formation of nP-collabs. The tau-R2 fragment with three pSers is in green and Na^+^ ions are shown as mauve van der Waals spheres. **(a)** One Na^+^ ion near pSer285 and a 2P-collab with pSer289 and pSer293, **(b)** 3P-collab involving several Na^+^ ions, **(c)** tripod involving pSer289, pSer293 and αI-Glu434.

### Impact of tau-R2 and its P-state on the CTT dynamics

Characterizing the dynamics of IDRs such as the CTTs at the residue level has always been challenging regardless of the method of investigation.^63-65^ The RMSF suffers from the need for a reference structure, and metrics like Protein Blocks are not so straightforward to use, as they require the user to first learn the structural alphabet in order to interpret the results.^66^ We recently defined Proteic Menger Curvatures (PMCs) as a simple way to describe backbone dynamics regardless of its structuration (or lack thereof).^38^ Curvature is an intuitive concept which also possesses the valuable property to relate to flexibility, as flexible objects display wide ranges of curvatures. This led us to define two additional relevant metrics, the Local Curvatures (LCs) and the Local Flexibilities (LFs), as the respective mean and standard deviation of the PMC for a given residue.

We applied the PMCs to the CTTs of the β-tubulin located on the N-terminal part of R2 and to the αI-CTT of each complex. In the absence of R2, the profile of the βIII-CTT PMCs shows that the tip of the CTT (residues Glu445 to Gly448) is mostly curved, whereas the base adopts an extended conformation as it stretches outward from the tubulin surface (see Figure 8b). When the βIII-CTT wraps around R2, following the mechanism described in Ref. 37 (see Figure 8c), the curvature shifts towards the base of the CTT, with two stable peaks around residues Glu442-Glu443 and Glu435-Met436. Although the CTT does not fold, its dynamics undergo a distinct shift upon wrapping around R2 (see Figure 8d). This transition from free to bound state of the tubulin CTTs upon tau-R2 binding was initially shown in ref. 37 (in figure 6) by monitoring the number of contacts formed between the CTT and the tubulin core and tau-R2. We report all PMC profiles of the CTT on the C-terminal side of R2 in Figure S6. In addition to the wrapping of the βI CTT identified for the second replica of the unphosphorylated state P000 in our previous study,^37^ we also witnessed it for the βI/αI/βI system in the P110 state and the βIII/αI/βIII system in the P001 state (see Figure S7 for the corresponding snapshots). Again, the curvature of the backbone is accentuated at the base of the CTT during the wrapping. These events show that wrapping can also take place around a phosphorylated fragment of Tau, despite the additional negative charges added by the phosphoserines. Since its occurrence rate is low, we cannot report on the rate of the wrapping phenomenon, but using the PMCs allows us to further characterize the CTT bound state at the residue level.

**Figure 8.**
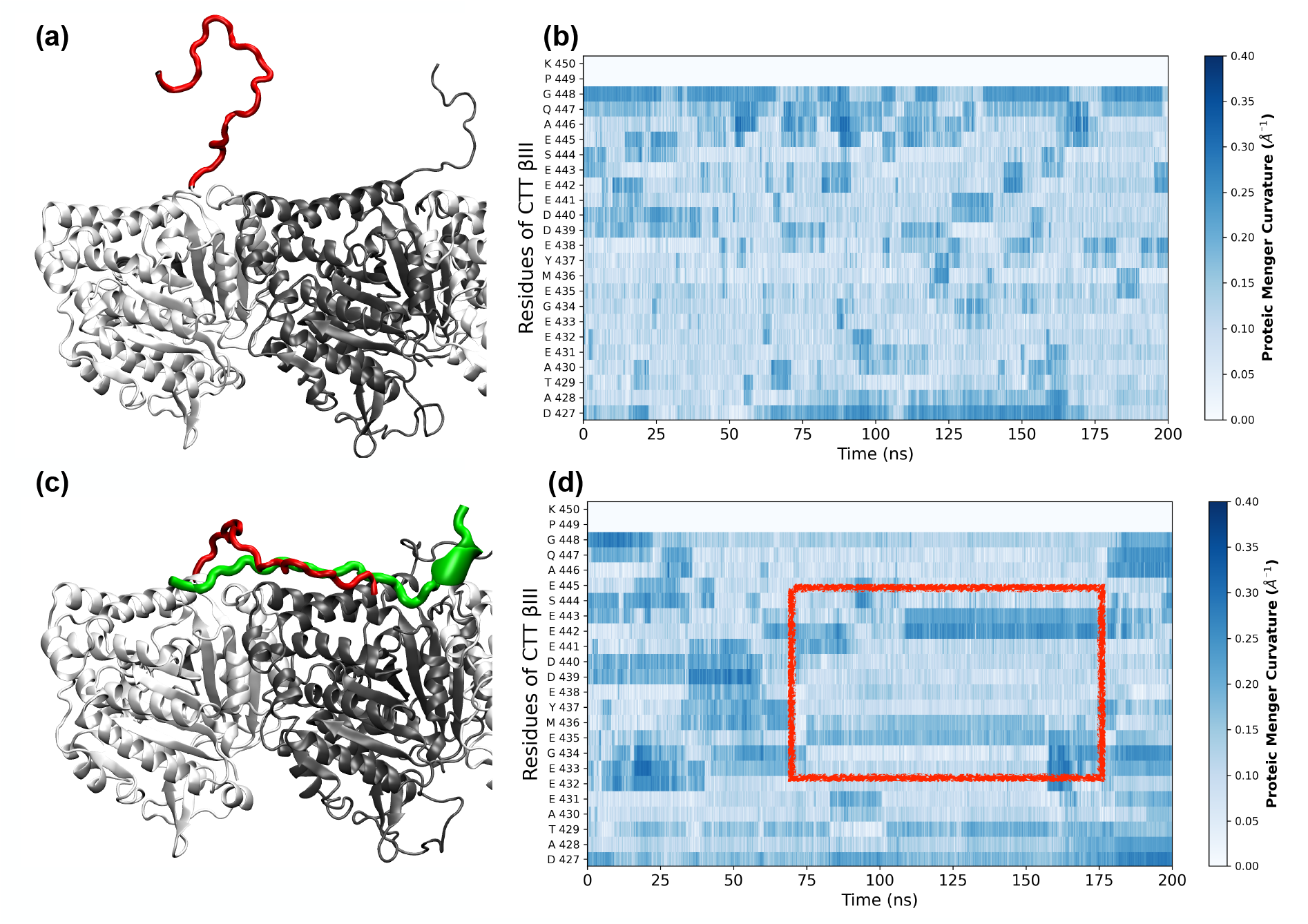
Left column, snapshots of the CTT for the first βIII-tubulin (in red) from the MD trajectories of the βIII/αI/βIII system (a) without tau-R2 or (c) with tau-R2 (in green) in the P000 state. Right column, local PMCs along time for the corresponding trajectories,(b) without tau, and (d) with tau. The red rectangle in the lower panel highlights the PMC increase due to the CTT wrapping around tau-R2.

R2-Ser285 interacts with the center of the α-tubulin surface, whereas R2-Ser289 and R2-Ser293 are located on the αβ interface. C-terminal residues R2-Lys294 to R2-Val300 are therefore in position to interact with the base of the αI-CTT. In the P000 state, residues R2-Lys294 to R2-Val300 interact transiently with residues αI-Ser439 to αI-Glu445 for the βI/αI/βI system but not for the βIII/αI/βIII one (see Figure 4b and c from Ref. 37). The opposite happens when Ser289 and Ser293 are phosphorylated, and we observe a significant increase of the contact time between R2-Lys294 and αI-Glu441. This increase is likely due to the strengthening of a salt bridge between R2-Lys294 and the phosphate group of pSer293, which allows αI-Glu441 to interact more easily with R2-Lys294 (see Figure 9 left panel). As a consequence, the LFs of the second half of the αI-CTT (residues αI-Glu446 to αI-Glu449) are reduced compared to the free CTT and the CTT in the non-phosphorylated system (see Figure 9 right panel).

**Figure 9.**
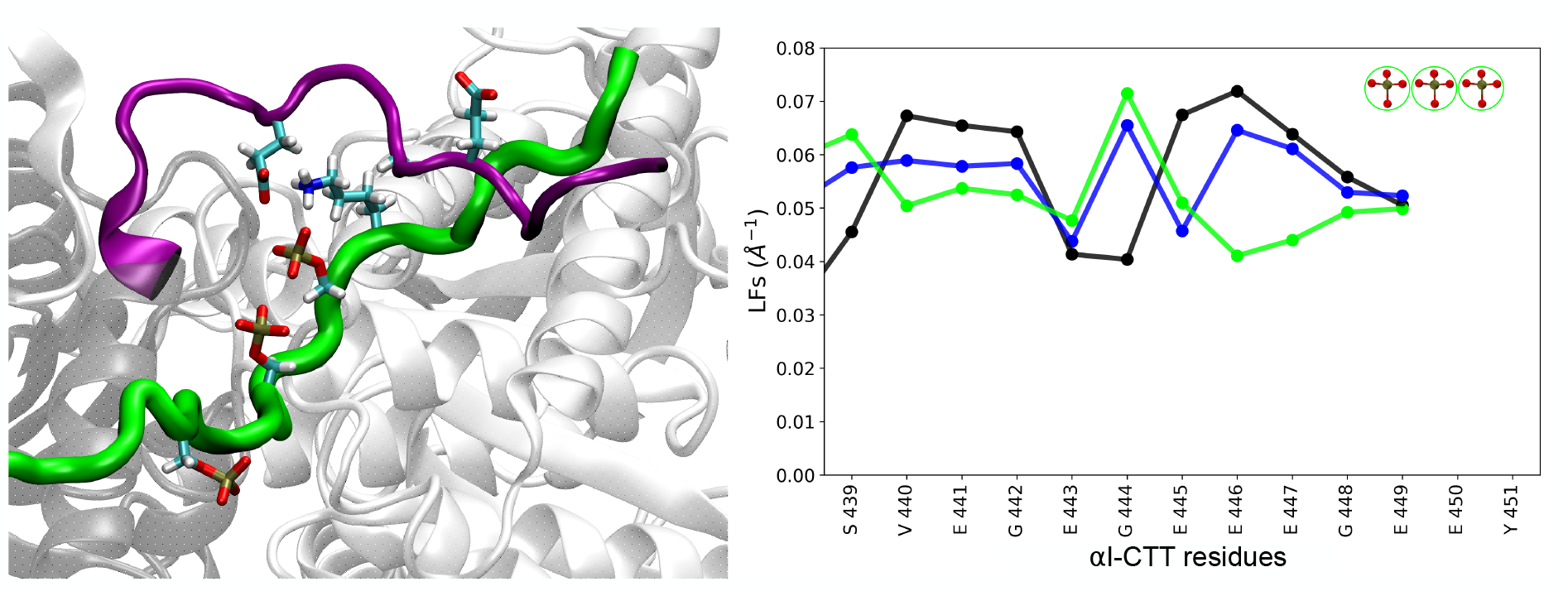
Left panel, snapshot of the CTT for the αI-tubulin (in purple) from the MD trajectories of the βIII/αI/βIII system with tau-R2 (in green) in the P111 state. Right panel, local flexibilities of the αI-CTT for the βIII/αI/βIII system without tau-R2 (black line) or with tau-R2 in the P000 (blue line) and P111 (green line) states.

## 4. Conclusions

The impact of phosphorylations on tau interaction with microtubules remains a complex problem, as, depending on its location along tau, this PTM can both disrupt^67-70^ or stabilize^71^ the tau/tubulin interface. In the present work, we used Molecular Dynamics simulations to investigate how the phosphorylation of serines 285, 289 and 293 impacts the interaction of the tau-R2 fragment with a tubulin trimer. As could be expected, and in agreement with earlier experimental and theoretical^62^ work, all eight possible P-states of tau-R2 present a decrease of their binding affinity for the tubulin surface. And this effect is more important when phosphorylating Ser289, which lies at the core of the strongly interacting region of tau-R2 that was defined by Brotzakis et al.^36^ One should also note that unlike Ser285 and Ser293, which can be phosphorylated in normal brains, Ser289 is the sole member of the serines R2 triad that is only phosphorylated in AD brains.^36, 72^ In addition, by including the tubulin CTTs in our simulation setup, we can show that their stabilizing effect on the tau/tubulin interface is still present in the phosphorylated states.

We solved the apparent contradiction between the low RMSF of phosphorylated tau on the MT surface and the large increase of the binding enthalpies upon phosphorylation by investigating the behavior of the Na^+^ counterions around the phosphate groups. The cations tend to bridge phosphate groups and contribute to stabilize the system, in a similar manner to what was observed in an earlier study modeling tau-R2 in solution.^38^ However, whether this bridging results from an artefactual behavior of CHARMM36m sodium counterions remains an open question. Finally, we show how tau-R2 phosphorylation also impacts the mobility and flexibility of the tubulin CTTs, as these can form new interactions with the phosphorylated R2 fragment. This in turn can impact the CTTs function as a MAP recruiters and regulators of motor proteins on the MT surface.

Further work will expand our approach by investigating additional phosphorylation spots along tau on a longer tubulin filament. This will help us to get rid of edge effects, which are likely to contribute to tau unbinding on its N-ter and C-ter termini. In addition, work on a larger system will improve the quality of our statistics regarding the mobility and conformational space covered by the disordered tubulin CTTs.

## Supporting Information Available

Additional data regarding the tubulin and tau sequences and modeling is available as a pdf file. All MD trajectories (without solvent) and their topologies for each simulation are deposited in Zenodo (https://zenodo.org/records/14888178).

## Acknowledgments

This work was supported by the ANR (MAGNETAU-ANR-21-CE29-0024) and by the “Initiative d’Excellence” program from the French State (Grant “DYNAMO”, ANR-11-LABX-0011-01 and grant CACSICE, ANR-11-EQPX-0008). Simulations were performed using the HPC resources from LBT/HPC and GENCI-TGCC (grant A0160714126).

## TOC image

For Table of Contents use only

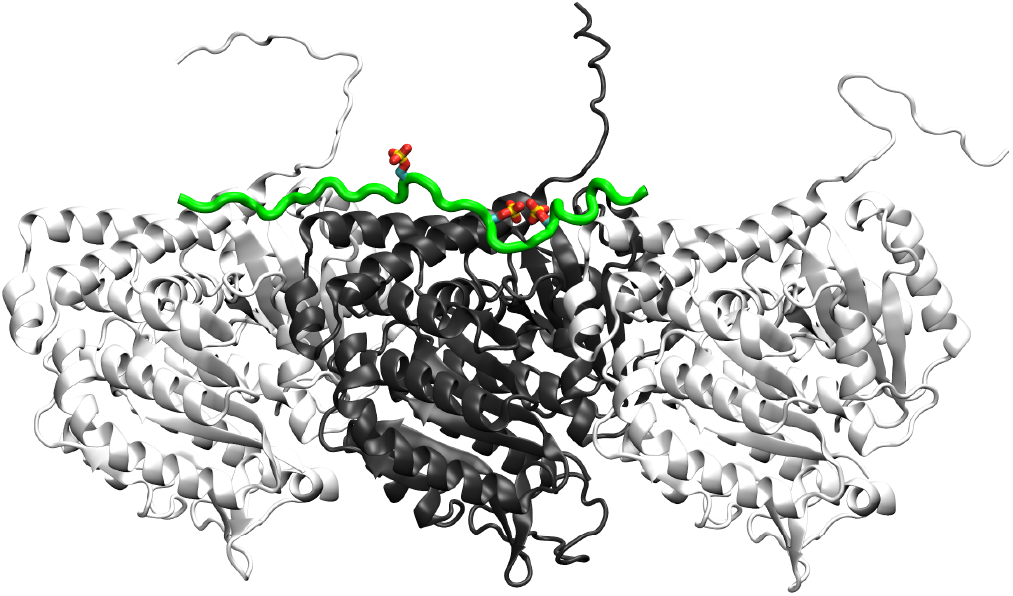

