## Supplementary information for "Impact of multiple phosphorylations on the tau-R2/tubulin interface"

*Jules Marien, Chantal Prevost and Sophie Sacquin-Mora**

Laboratoire de Biochimie Théorique, Université Paris-Cité, CNRS,

13 rue Pierre et Marie Curie, Paris 75005, France

**Table S1:** Homology percentage and sequence alignments between the template sequence (from pdb 6CVN) and the sheep and human tubulins used in the study.

| | αI (TUBAIA) | βI (TUBB) | βIII (TUBB3) |
|---|---|---|---|
| 6CVN (α subunit) | 99,5 % | | |
| 6CVN (β subunit) | | 97,9 % | 94,4 % |

```
alpha1_core_sheep      MRECISIHVGQAGVQIGNACWELYCLEHGIQPDGQMPSDKTIGGGDDSFNTFFSETGAGK
alpha1_6CVN            MRECISIHVGQAGVQIGNACWELYCLEHGIQPDGQMPSDKTIGGGDDSFNTFFSETGAGK
                       ************************************************************

alpha1_core_sheep      HVPRAVFVDLEPTVIDEVRTGTYRQLFHPEQLITGKEDAANNYARGHYTIGKEIIDLVLD
alpha1_6CVN            HVPRAVFVDLEPTVIDEVRTGTYRQLFHPEQLITGKEDAANNYARGHYTIGKEIIDLVLD
                       ************************************************************

alpha1_core_sheep      RIRKLADQCTGLQGFLVFHSFGGGTGSGFTSLLMERLSVDYGKKSKLEFSIYPAPQVSTA
alpha1_6CVN            RIRKLADQCTGLQGFLVFHSFGGGTGSGFTSLLMERLSVDYGKKSKLEFSIYPAPQVSTA
                       ************************************************************

alpha1_core_sheep      VVEPYNSILTTHTTLEHSDCAFMVDNEAIYDICRRNLDIERPTYTNLNRLIGQIVSSITA
alpha1_6CVN            VVEPYNSILTTHTTLEHSDCAFMVDNEAIYDICRRNLDIERPTYTNLNRLISQIVSSITA
                       ***************************************************.********

alpha1_core_sheep      SLRFDGALNVDLTEFQTNLVPYPRIHFPLATYAPVISAEKAYHEQLSVAEITNACFEPAN
alpha1_6CVN            SLRFDGALNVDLTEFQTNLVPYPRIHFPLATYAPVISAEKAYHEQLSVAEITNACFEPAN
                       ************************************************************

alpha1_core_sheep      QMVKCDPRHGKYMACCLLYRGDVVPKDVNAAIATIKTKRTIQFVDWCPTGFKVGINYQPP
alpha1_6CVN            QMVKCDPRHGKYMACCLLYRGDVVPKDVNAAIATIKTKRSIQFVDWCPTGFKVGINYQPP
                       ***************************************.********************

alpha1_core_sheep      TVVPGGDLAKVQRAVCMLSNTTAIAEAWARLDHKFDLMYAKRAFVHWYVGEGMEEGEFSE
alpha1_6CVN            TVVPGGDLAKVQRAVCMLSNTTAIAEAWARLDHKFDLMYAKRAFVHWYVGEGMEEGEFSE
                       ************************************************************

alpha1_core_sheep      AREDMAALEKDYEEVGVD
alpha1_6CVN            AREDMAALEKDYEEVGVD
                       ******************
```

```
beta1_core_sheep      MREIVHIQAGQCGNQIGAKFWEVISDEHGIDPTGTYHGDSDLQLDRISVYYNEATGGKYV
beta1_6CVN            MREIVHIQAGQCGNQIGAKFWEVISDEHGIDPTGSYHGDSDLQLERINVYYNEAAGNKYV
                      **********************************:*********:**.******:*.***

beta1_core_sheep      PRAILVDLEPGTMDSVRSGPFGQIFRPDNFVFGQSGAGNNWAKGHYTEGAELVDSVLDVV
beta1_6CVN            PRAILVDLEPGTMDSVRSGPFGQIFRPDNFVFGQSGAGNNWAKGHYTEGAELVDSVLDVV
                      ************************************************************

beta1_core_sheep      RKEAESCDCLQGFQLTHSLGGGTGSGMGTLLISKIREEYPDRIMNTFSVVPSPKVSDTVV
beta1_6CVN            RKESESCDCLQGFQLTHSLGGGTGSGMGTLLISKIREEYPDRIMNTFSVVPSPKVSDTVV
                      ***:********************************************************

beta1_core_sheep      EPYNATLSVHQLVENTDETYCIDNEALYDICFRTLKLTTPTYGDLNHLVSATMSGVTTCL
beta1_6CVN            EPYNATLSVHQLVENTDETYCIDNEALYDICFRTLKLTTPTYGDLNHLVSATMSGVTTCL
                      ************************************************************

beta1_core_sheep      RFPGQLNADLRKLAVNMVPFPRLHFFMPGFAPLTSRGSQQYRALTVPELTQQVFDAKNMM
beta1_6CVN            RFPGQLNADLRKLAVNMVPFPRLHFFMPGFAPLTSRGSQQYRALTVPELTQQMFDAKNMM
                      ****************************************************:*******

beta1_core_sheep      AACDPRHGRYLTVAAVFRGRMSMKEVDEQMLNVQNKNSSYFVEWIPNNVKTAVCDIPPRG
beta1_6CVN            AACDPRHGRYLTVAAVFRGRMSMKEVDEQMLNVQNKNSSYFVEWIPNNVKTAVCDIPPRG
                      ************************************************************

beta1_core_sheep      LKMAVTFIGNSTAIQELFKRISEQFTAMFRRKAFLHWYTGEGMDEMEFTEAESNMNDLVS
beta1_6CVN            LKMSATFIGNSTAIQELFKRISEQFTAMFRRKAFLHWYTGEGMDEMEFTEAESNMNDLVS
                      ***:.*******************************************************

beta1_core_sheep      EYQQYQ
beta1_6CVN            EYQQYQ
                      ******


beta3_core_human      MREIVHIQAGQCGNQIGAKFWEVISDEHGIDPSGNYVGDSDLQLERISVYYNEASSHKYV
beta1_6CVN            MREIVHIQAGQCGNQIGAKFWEVISDEHGIDPTGSYHGDSDLQLERINVYYNEAAGNKYV
                      ********************************:*.* **********.******:.:***

beta3_core_human      PRAILVDLEPGTMDSVRSGAFGHLFRPDNFIFGQSGAGNNWAKGHYTEGAELVDSVLDVV
beta1_6CVN            PRAILVDLEPGTMDSVRSGPFGQIFRPDNFVFGQSGAGNNWAKGHYTEGAELVDSVLDVV
                      ******************* **::******:*****************************

beta3_core_human      RKECENCDCLQGFQLTHSLGGGTGSGMGTLLISKVREEYPDRIMNTFSVVPSPKVSDTVV
beta1_6CVN            RKESESCDCLQGFQLTHSLGGGTGSGMGTLLISKIREEYPDRIMNTFSVVPSPKVSDTVV
                      ***.*.****************************:*************************

beta3_core_human      EPYNATLSIHQLVENTDETYCIDNEALYDICFRTLKLATPTYGDLNHLVSATMSGVTTSL
beta1_6CVN            EPYNATLSVHQLVENTDETYCIDNEALYDICFRTLKLTTPTYGDLNHLVSATMSGVTTCL
                      ********:****************************:********************.*

beta3_core_human      RFPGQLNADLRKLAVNMVPFPRLHFFMPGFAPLTARGSQQYRALTVPELTQQMFDAKNMM
beta1_6CVN            RFPGQLNADLRKLAVNMVPFPRLHFFMPGFAPLTSRGSQQYRALTVPELTQQMFDAKNMM
                      **********************************:*************************

beta3_core_human      AACDPRHGRYLTVATVFRGRMSMKEVDEQMLAIQSKNSSYFVEWIPNNVKVAVCDIPPRG
beta1_6CVN            AACDPRHGRYLTVAAVFRGRMSMKEVDEQMLNVQNKNSSYFVEWIPNNVKTAVCDIPPRG
                      **************:**************** :*.***************.*********

beta3_core_human      LKMSSTFIGNSTAIQELFKRISEQFTAMFRRKAFLHWYTGEGMDEMEFTEAESNMNDLVS
beta1_6CVN            LKMSATFIGNSTAIQELFKRISEQFTAMFRRKAFLHWYTGEGMDEMEFTEAESNMNDLVS
                      ****:*******************************************************

beta3_core_human      EYQQYQ
beta1_6CVN            EYQQYQ
                      ******
```

**Table S2:** Aligned sequences of the tubulin C-terminal tails

| Isotype | Sequence | Length |
|---|---|---|
| αI (D0vWZ0) | SVEGEGEEEGEEY | 13 |
| βI (W5PPT6) | DATAEEEE--DFGEEAEEEA | 18 |
| βIII (Q13509) | DATAEEEG--EMYEDDEEESEAQGPK | 24 |

**Table S3:** Summary of the MD simulations performed in the study (for each system/P-state combination, we ran three replicas of 200 ns).

composition_system

| System | P-state | Protein atoms | Sodium ions | Chloride ions | Water molecules |
|---|---|---|---|---|---|
| **no CTTs** | 0 0 0 | 20377 | 308 | 267 | 93635 |
| **no CTTs** | 0 0 1 | 20380 | 310 | 267 | 93662 |
| **no CTTs** | 0 1 0 | 20380 | 310 | 267 | 93653 |
| **no CTTs** | 0 1 1 | 20383 | 312 | 267 | 93645 |
| **no CTTs** | 1 0 0 | 20380 | 310 | 267 | 93641 |
| **no CTTs** | 1 0 1 | 20383 | 312 | 267 | 93617 |
| **no CTTs** | 1 1 0 | 20383 | 312 | 267 | 93603 |
| **no CTTs** | 1 1 1 | 20386 | 314 | 267 | 93608 |
| **βI/αI/βI** | 0 0 0 | 21031 | 336 | 266 | 93216 |
| **βI/αI/βI** | 0 0 1 | 21034 | 338 | 266 | 93248 |
| **βI/αI/βI** | 0 1 0 | 21034 | 338 | 266 | 93229 |
| **βI/αI/βI** | 0 1 1 | 21037 | 340 | 266 | 93196 |
| **βI/αI/βI** | 1 0 0 | 21034 | 338 | 266 | 93213 |
| **βI/αI/βI** | 1 0 1 | 21037 | 340 | 266 | 93199 |
| **βI/αI/βI** | 1 1 0 | 21037 | 340 | 266 | 93244 |
| **βI/αI/βI** | 1 1 1 | 21040 | 342 | 266 | 93220 |
| **βIII/αI/βIII** | 0 0 0 | 21227 | 426 | 356 | 125253 |
| **βIII/αI/βIII** | 0 0 1 | 21230 | 428 | 356 | 125223 |
| **βIII/αI/βIII** | 0 1 0 | 21230 | 428 | 356 | 125229 |
| **βIII/αI/βIII** | 0 1 1 | 21233 | 430 | 356 | 125196 |
| **βIII/αI/βIII** | 1 0 0 | 21230 | 428 | 356 | 125229 |
| **βIII/αI/βIII** | 1 0 1 | 21233 | 430 | 356 | 125211 |
| **βIII/αI/βIII** | 1 1 0 | 21233 | 430 | 356 | 125223 |
| **βIII/αI/βIII** | 1 1 1 | 21236 | 432 | 356 | 125215 |

**Table S4:** Average binding enthalpies (in kcal.mol$^{-1}$), with the different contributions, between tau-R2 and the tubulin heterotrimer. Calculations were performed with a stride of 10 frames on the last 150 ns of trajectory, and the results from the three replicas were combined for each system.

| System | p-state | Total | Elec | Vdw | PB | SASA |
|---|---|---|---|---|---|---|
| **no CTTs** | 0 0 0 | -55.0 +/- 18.6 | -2236.5 +/- 206.1 | -73.3 +/- 12.3 | 2266.0 +/- 205.4 | -11.1 +/- 1.5 |
| **βI/αI/βI** | 0 0 0 | -65.8 +/- 23.1 | -3131.7 +/- 237.1 | -78.9 +/- 18.2 | 3157.3 +/- 235.5 | -12.4 +/- 2.6 |
| **βIII/αI/βIII** | 0 0 0 | -73.2 +/- 21.5 | -3198.5 +/- 250.3 | -73.8 +/- 15.6 | 3211.0 +/- 252.2 | -11.9 +/- 2.3 |
| **no CTTs** | 0 0 1 | -24.9 +/- 19.9 | -1374.1 +/- 218.7 | -49.3 +/- 13.2 | 1407.0 +/- 211.0 | -8.5 +/- 1.6 |
| **βI/αI/βI** | 0 0 1 | -50.3 +/- 29.6 | -1687.4 +/- 217.8 | -63.0 +/- 20.7 | 1711.0 +/- 204.7 | -10.8 +/- 1.6 |
| **βIII/αI/βIII** | 0 0 1 | -55.6 +/- 24.0 | -1896.7 +/- 237.4 | -76.9 +/- 23.9 | 1929.9 +/- 236.3 | -11.9 +/- 2.8 |
| **no CTTs** | 0 1 0 | -0.7 +/- 25.6 | -1261.7 +/- 177.8 | -68.6 +/- 14.1 | 1340.4 +/- 158.5 | -10.7 +/- 1.3 |
| **βI/αI/βI** | 0 1 0 | -32.0 +/- 19.4 | -1577.8 +/- 215.2 | -63.2 +/- 23.8 | 1619.4 +/- 222.6 | -10.4 +/- 3.5 |
| **βIII/αI/βIII** | 0 1 0 | -25.5 +/- 21.6 | -1715.3 +/- 136.4 | -81.5 +/- 10.0 | 1783.6 +/- 128.3 | -12.4 +/- 1.1 |
| **no CTTs** | 1 0 0 | -40.2 +/- 23.5 | -1472.0 +/- 245.2 | -65.5 +/- 11.4 | 1507.6 +/- 231.3 | -10.3 +/- 1.7 |
| **βI/αI/βI** | 1 0 0 | -55.7 +/- 18.3 | -1849.1 +/- 182.5 | -75.4 +/- 16.3 | 1880.5 +/- 178.9 | -11.8 +/- 2.0 |
| **βIII/αI/βIII** | 1 0 0 | -51.0 +/- 21.6 | -1765.7 +/- 260.4 | -81.2 +/- 16.5 | 1808.3 +/- 252.6 | -12.4 +/- 1.8 |
| **no CTTs** | 0 1 1 | -9.6 +/- 32.7 | -508.6 +/- 230.7 | -53.1 +/- 22.9 | 561.7 +/- 214.1 | -9.6 +/- 2.3 |
| **βI/αI/βI** | 0 1 1 | 25.1 +/- 50.8 | -273.8 +/- 279.2 | -74.1 +/- 14.9 | 384.8 +/- 245.0 | -11.8 +/- 1.7 |
| **βIII/αI/βIII** | 0 1 1 | -19.6 +/- 26.1 | -537.9 +/- 220.3 | -79.3 +/- 12.1 | 611.1 +/- 198.9 | -13.5 +/- 1.1 |
| **no CTTs** | 1 0 1 | -20.1 +/- 23.5 | -438.8 +/- 222.9 | -59.7 +/- 11.4 | 488.3 +/- 222.7 | -9.9 +/- 1.6 |
| **βI/αI/βI** | 1 0 1 | -40.2 +/- 21.1 | -509.9 +/- 169.2 | -63.4 +/- 22.6 | 544.1 +/- 166.9 | -11.0 +/- 3.4 |
| **βIII/αI/βIII** | 1 0 1 | -39.4 +/- 23.3 | -458.1 +/- 202.2 | -82.3 +/- 14.5 | 513.4 +/- 189.5 | -12.4 +/- 1.7 |
| **no CTTs** | 1 1 0 | 12.2 +/- 27.4 | -442.1 +/- 167.7 | -66.0 +/- 12.6 | 531.0 +/- 166.0 | -10.7 +/- 1.6 |
| **βI/αI/βI** | 1 1 0 | -7.7 +/- 44.3 | -454.5 +/- 301.7 | -87.6 +/- 14.7 | 548.4 +/- 275.9 | -14.0 +/- 2.1 |
| **βIII/αI/βIII** | 1 1 0 | -13.2 +/- 20.9 | -393.9 +/- 211.3 | -75.4 +/- 22.6 | 467.9 +/- 207.4 | -11.8 +/- 2.4 |
| **no CTTs** | 1 1 1 | 36.2 +/- 28.0 | 277.4 +/- 232.7 | -49.9 +/- 12.5 | -182.2 +/- 216.9 | -9.1 +/- 1.3 |
| **βI/αI/βI** | 1 1 1 | -9.4 +/- 37.9 | 805.6 +/- 342.3 | -64.0 +/- 14.3 | -740.3 +/- 321.3 | -10.8 +/- 1.6 |
| **βIII/αI/βIII** | 1 1 1 | 37.7 +/- 25.8 | 1170.2 +/- 242.9 | -64.0 +/- 19.4 | -1057.4 +/- 229.4 | -11.1 +/- 2.6 |

**Figure S1:** Variations RMSD of the tau-R2 peptide as a function of time. Each panel shows the values for the three replicas for a given phosphorylation state. a) System with no CTTs.

(A)

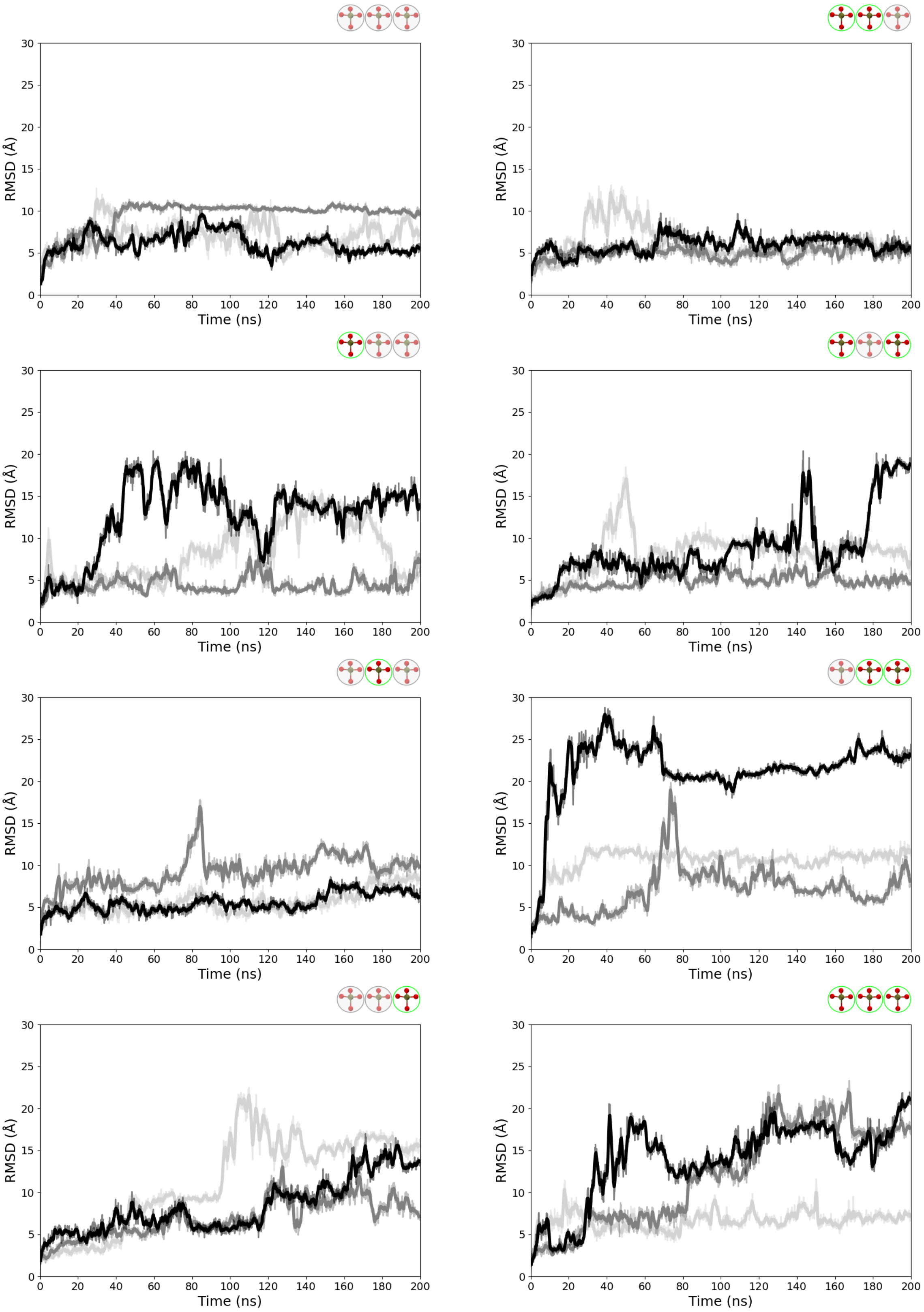

**Figure S1:** Variations RMSD of the tau-R2 peptide as a function of time. Each panel shows the values for the three replicas for a given phosphorylation state. b) system with the βI/αI/βI isotype.

(B)

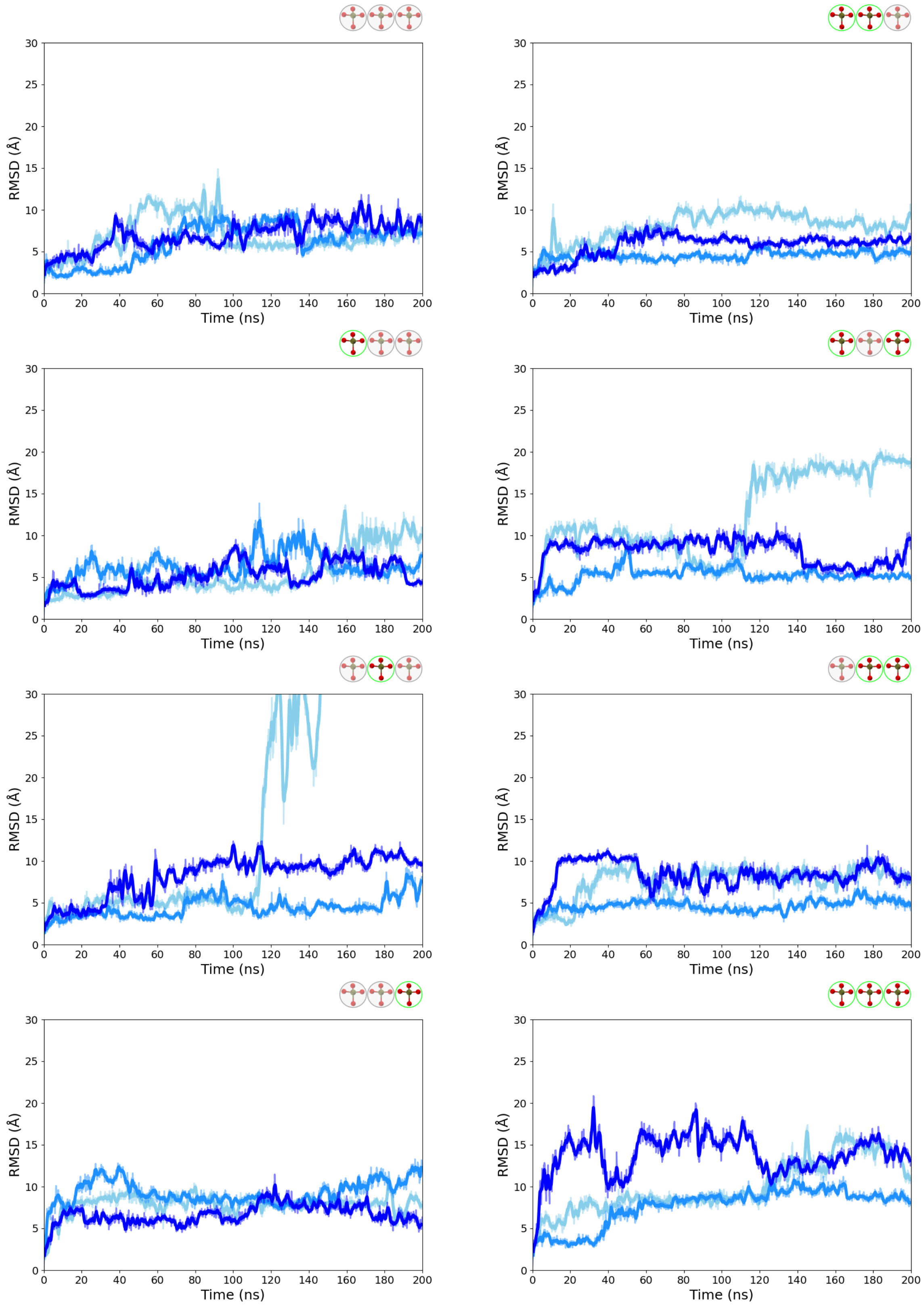

**Figure S1:** Variations RMSD of the tau-R2 peptide as a function of time. Each panel shows the values for the three replicas for a given phosphorylation state. c) System with the βIII/αI/βIII isotype.

(C)

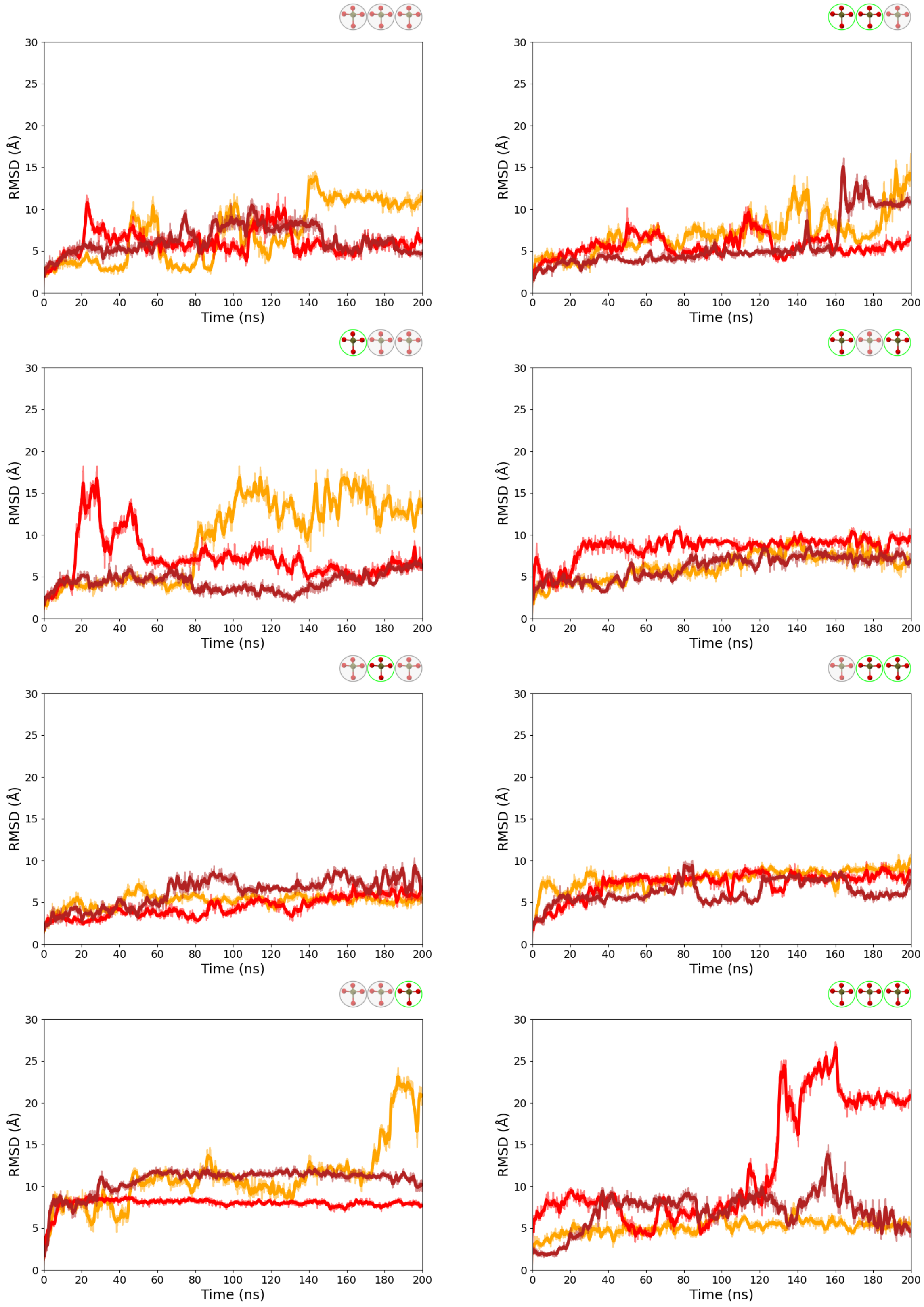

**Figure S2:** Variations in the tau-R2/tubulin contact maps for all P-states compared to the P000 states. The tubulin CTTs for the βI/αI/βI and βIII/αI/βIII isotypes (central and lower panels) are delimited by a purple frame, while the phosphorylated serines (Ser 285, 289 and 293) of tau-R2 are highlighted by green frames. Blue areas highlight a contact loss upon phosphorylation, while the red areas highlight a contact increase. (a) P001, (b) P010, (c) P100

(a)

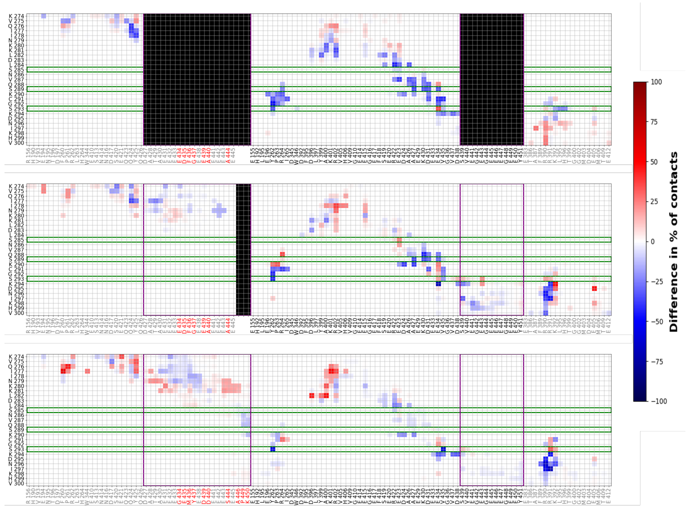


(b)

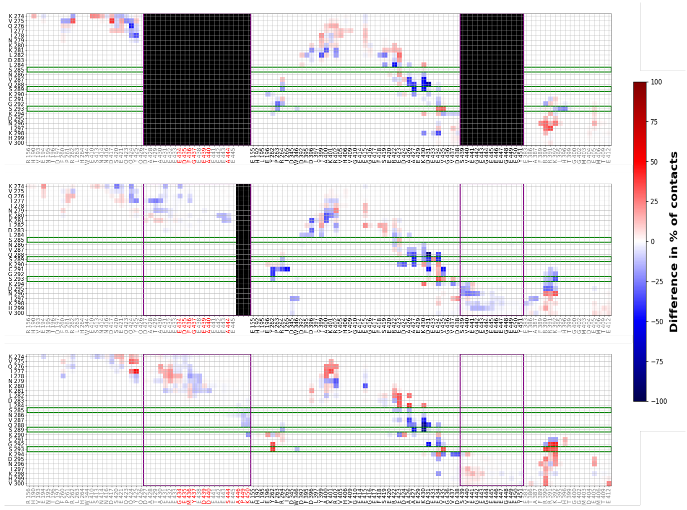


(c)

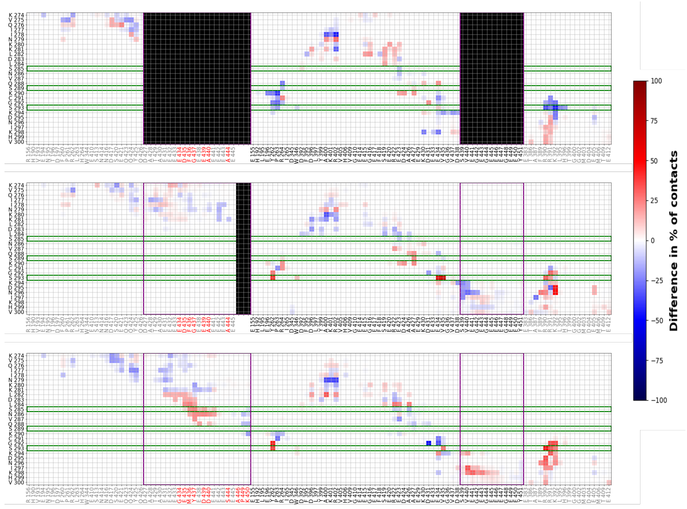

**Figure S2 (continued):** (d) P011, (e) P101, (f) P110

(d)

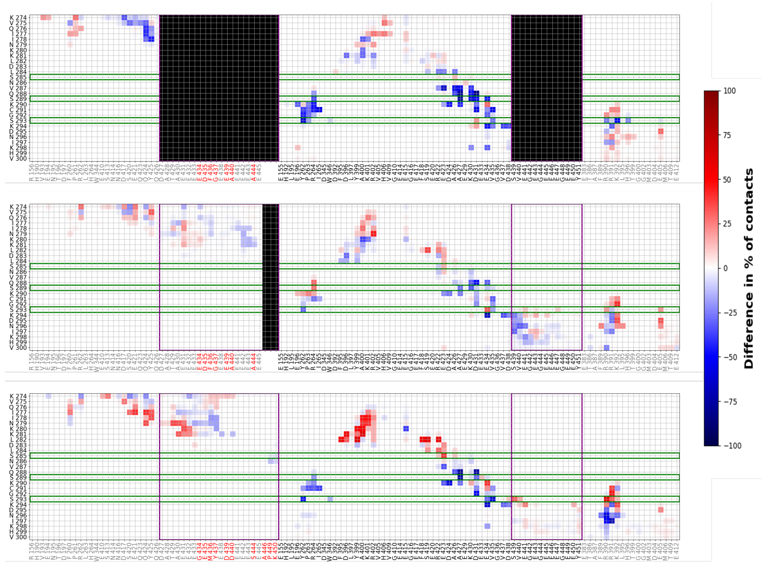

(e)

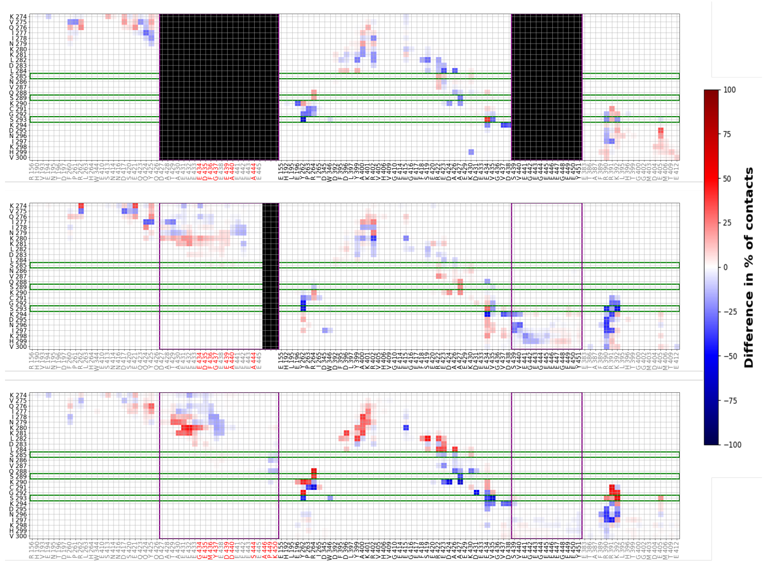

(f)

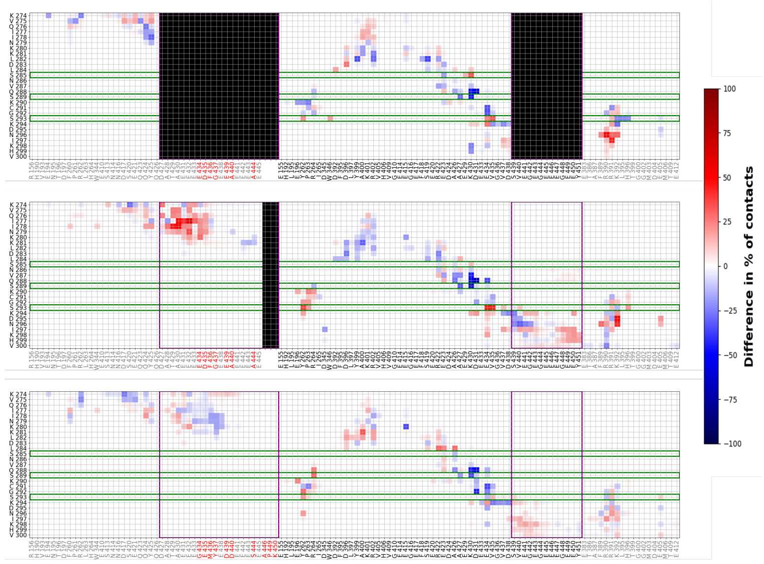

**Figure S3:** Number of R2/tubulin contacts as a function of the native contacts loss (1-Fnat) in all Pstates (each line displays the three replicas for a given Pstate). a) system without tubulin CTTs

(A)

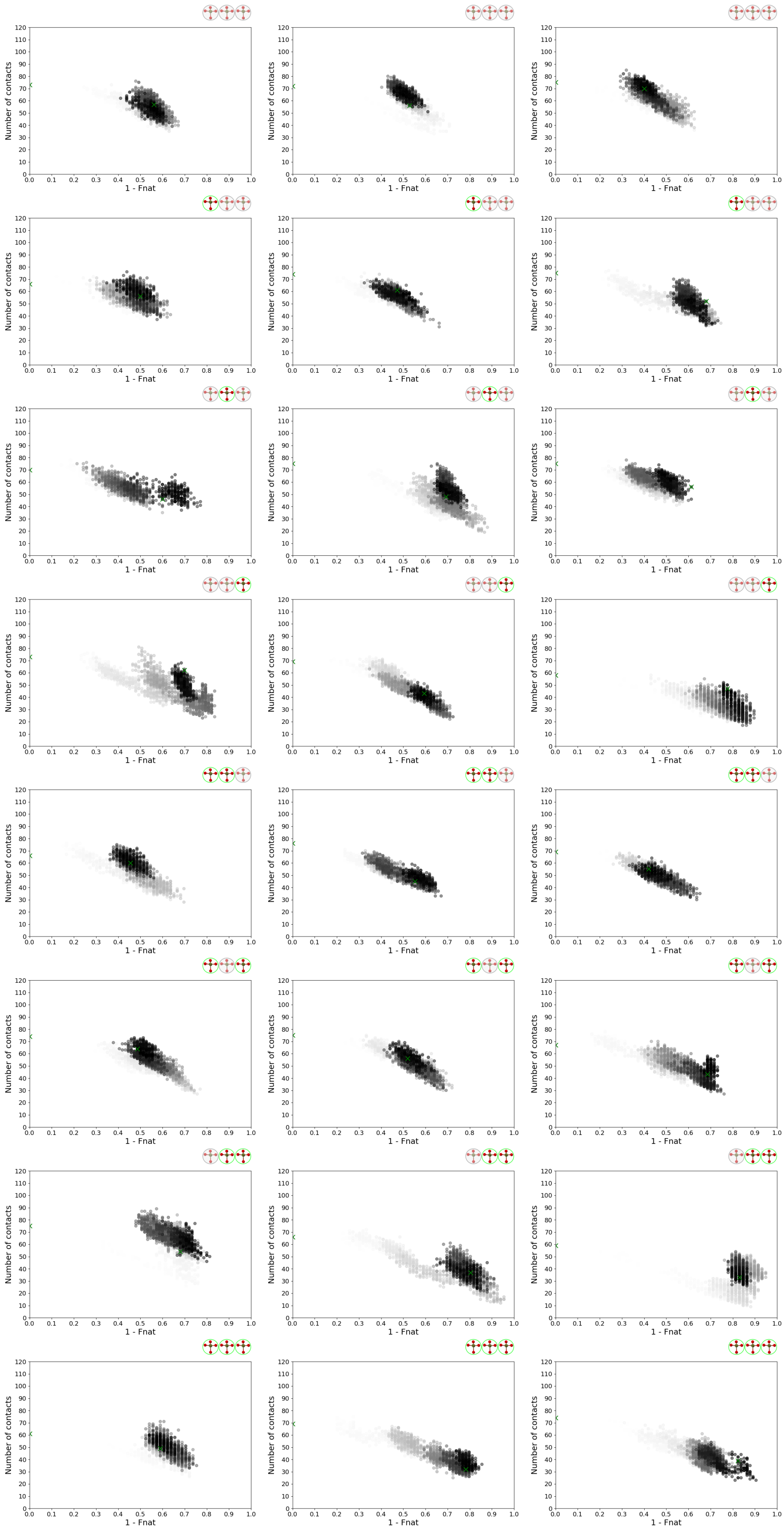

**Figure S3:** Number of R2/tubulin contacts as a function of the native contacts loss (1-Fnat) in all Pstates (each line displays the three replicas for a given Pstate). b) system with the βI/αI/βI isotype

(B)

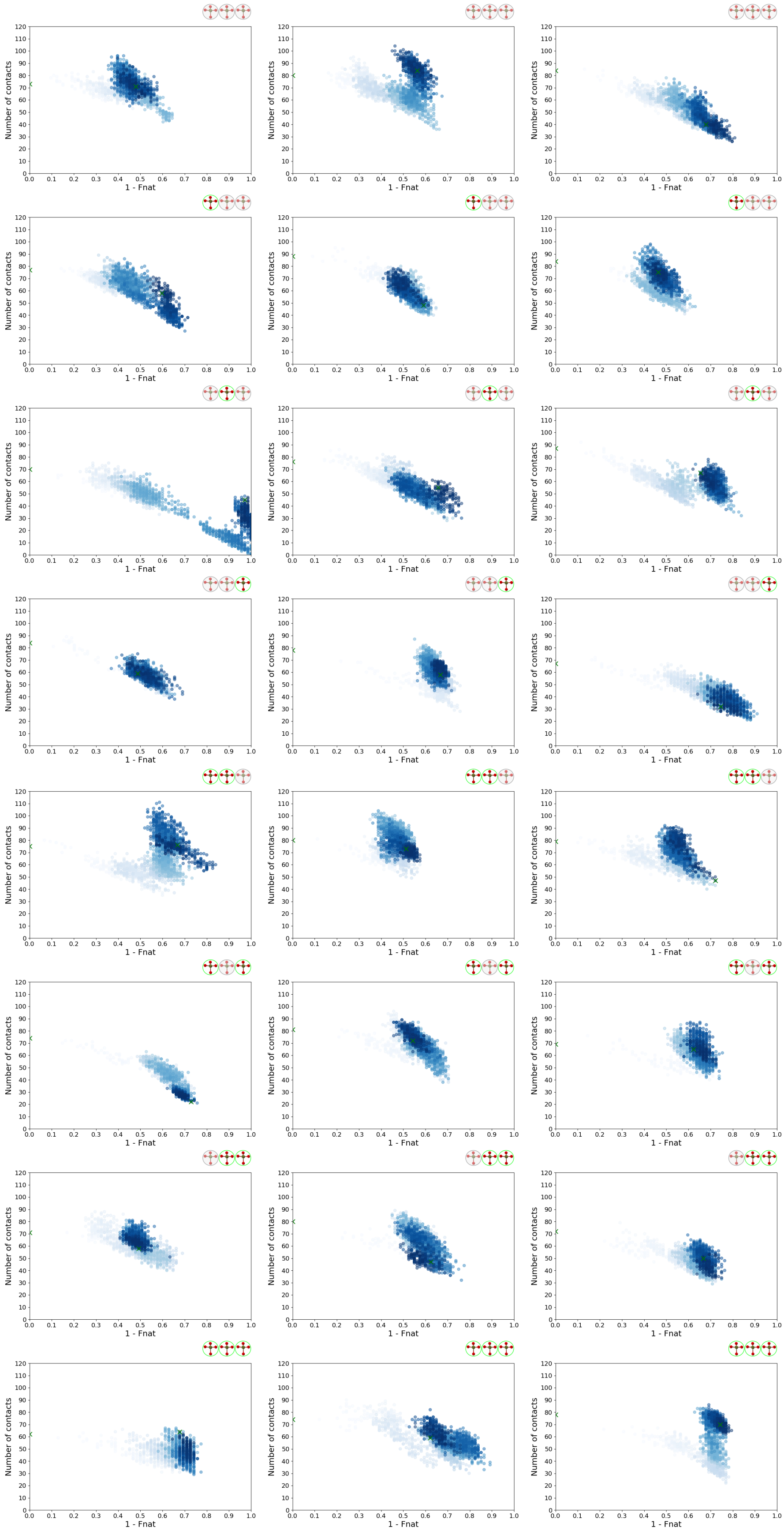

**Figure S3:** Number of R2/tubulin contacts as a function of the native contacts loss (1-Fnat) in all Pstates (each line displays the three replicas for a given Pstate). c) system with the βIII/αI/βIII isotype

(C)

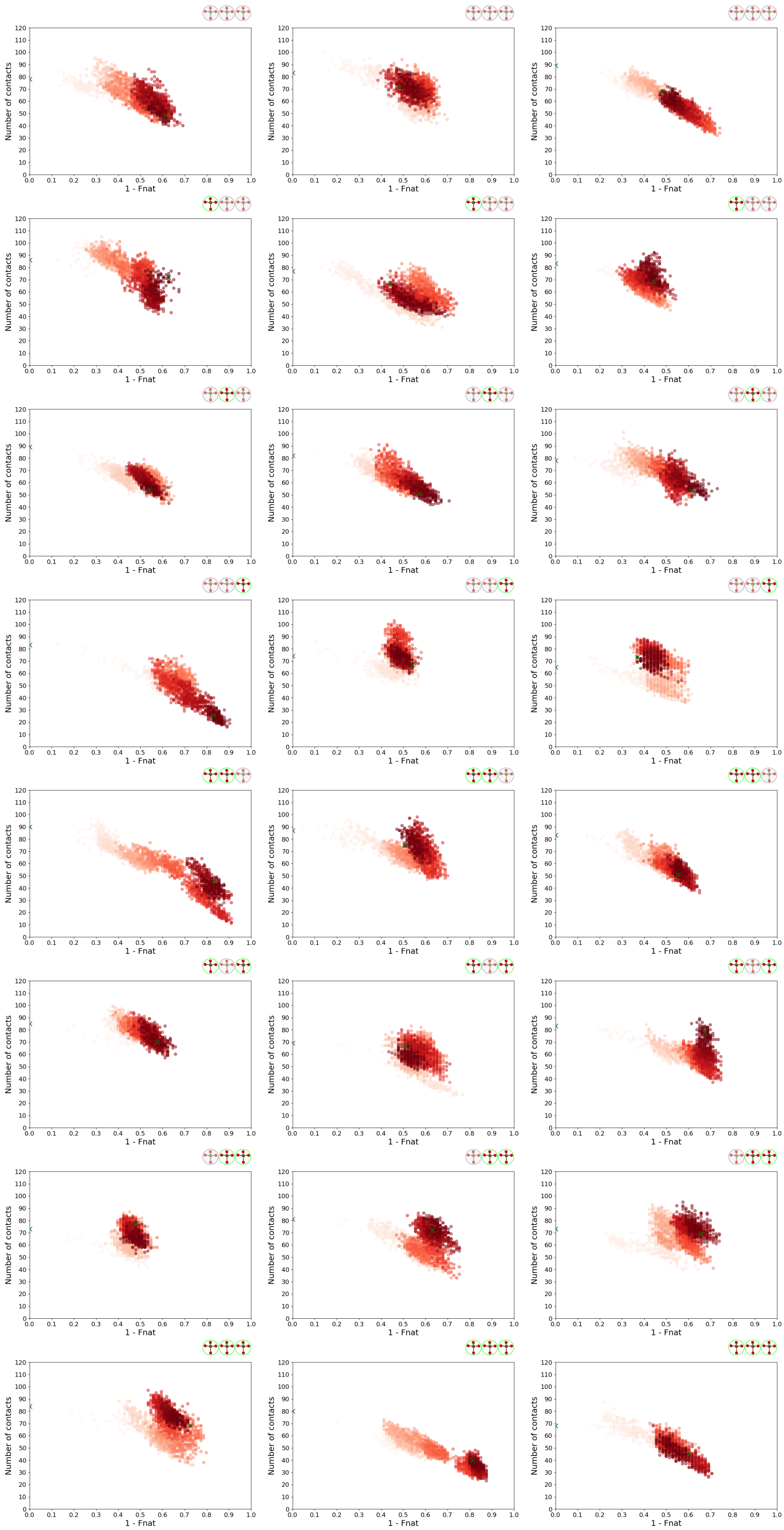

**Figure S4:** Different contributions (as listed in Table S4) to the binding enthalpies between tau-R2 and the tubulin heterotrimer for all the P-states in the three systems. (a) Electrostatic, (b) Van der Waals, (c) Poisson-Boltzmann, (d) Solvent accessible surface area.

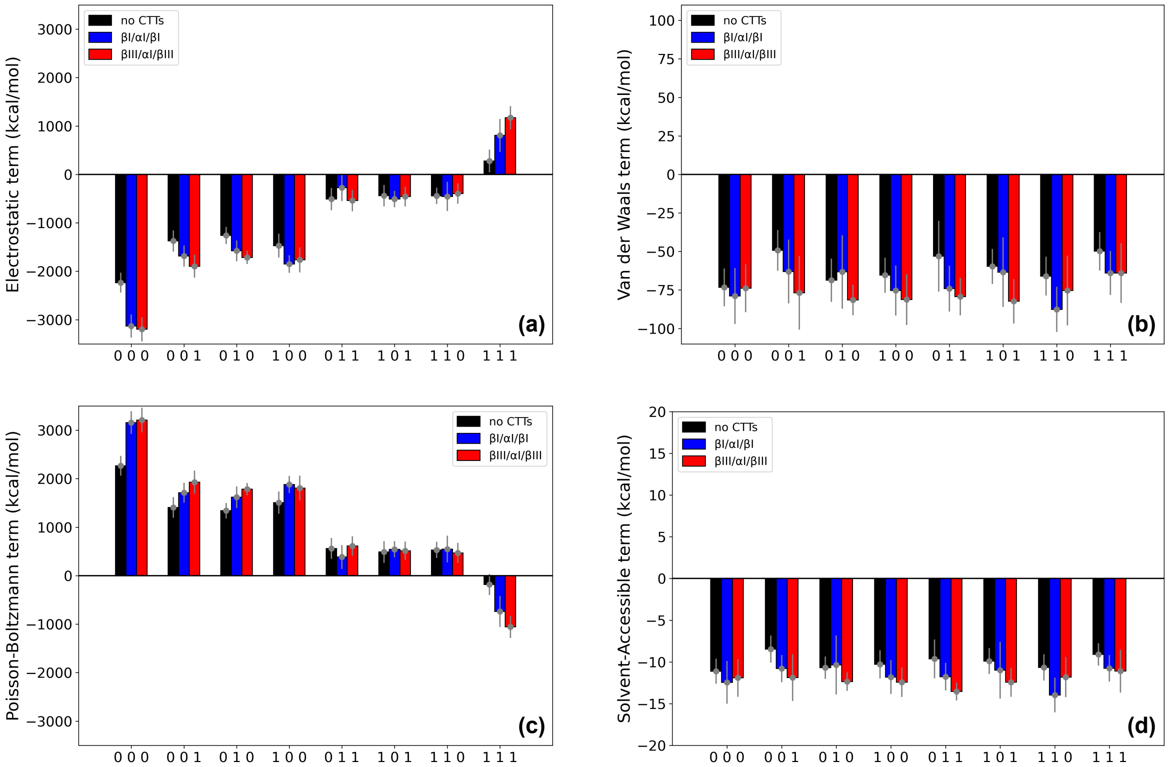

**Figure S5:** Monitoring the presence of $Na^{+}$ counterions in the vicinity of the R2 serines along the MD trajectories for each replica of the βI/αI/βI and βIII/αI/βIII system. Each horizontal line corresponds to one individual cation. Light blue dots indicate that the cation is less than 4 Å away from a P atom, dark blue indicate proximity to two P atoms (2P-collabs) and red proximity to the three P atoms (3P-collabs). (a) βIII/αI/βIII isotypes (P-states not shown in the manuscript main body), (b) βI/αI/βI (all P-states)

(a)

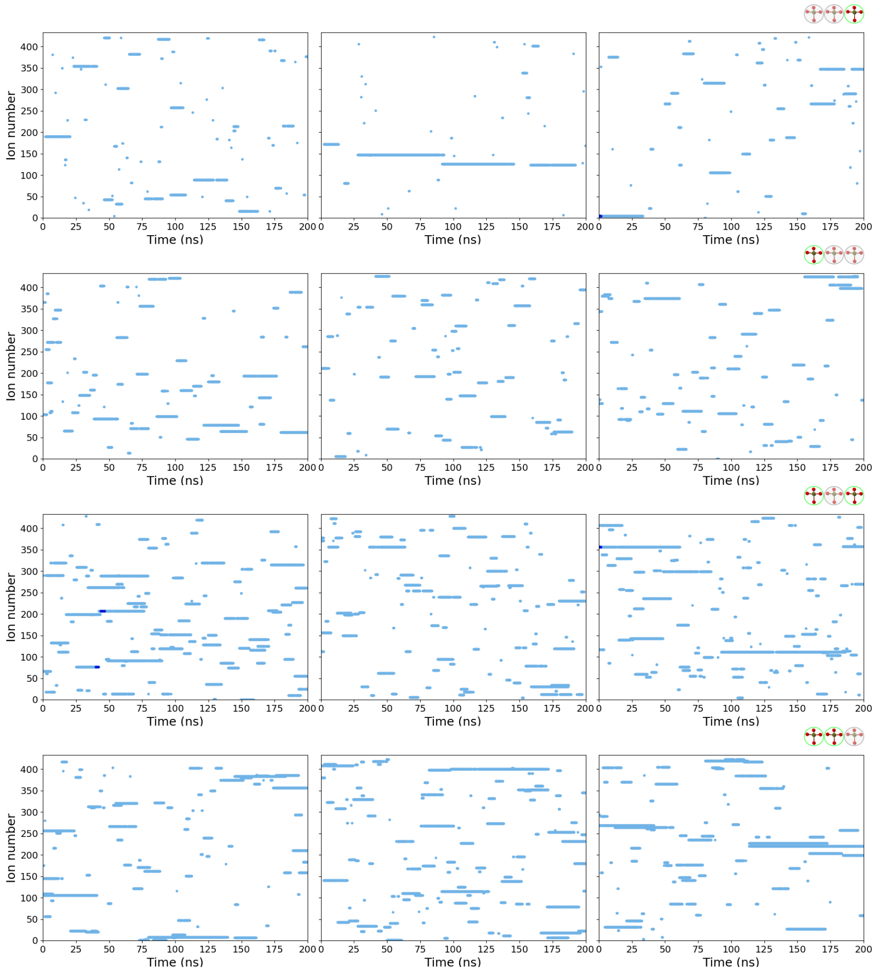

(b)

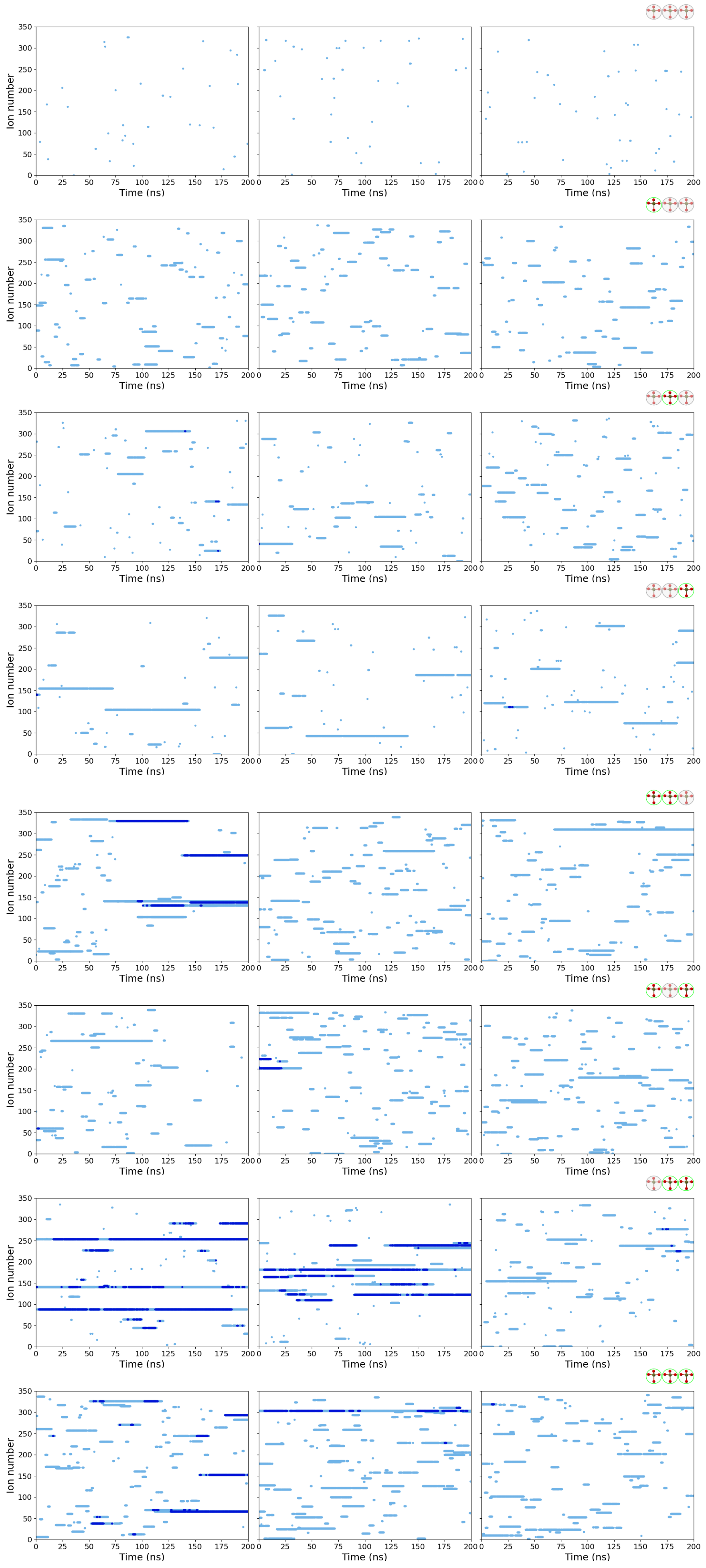

**Figure S6:** Local PMCs along time for all trajectories (8 Pstates with 3 replicas each).

a) βI/αI/βI system.

(A)

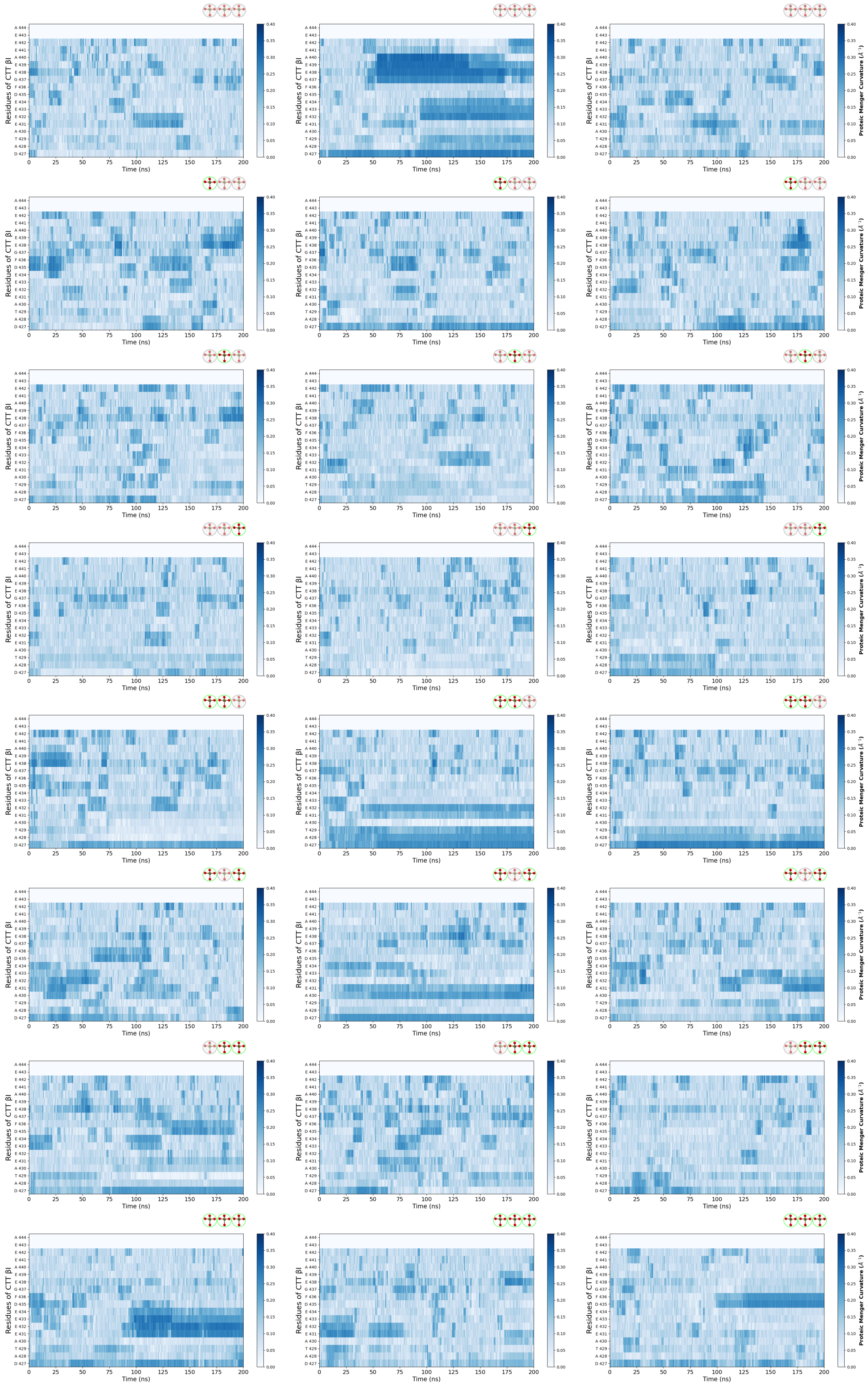

**Figure S6:** Local PMCs along time for all trajectories (8 Pstates with 3 replicas each).

b) βIII/αI/βIII system.

(B)

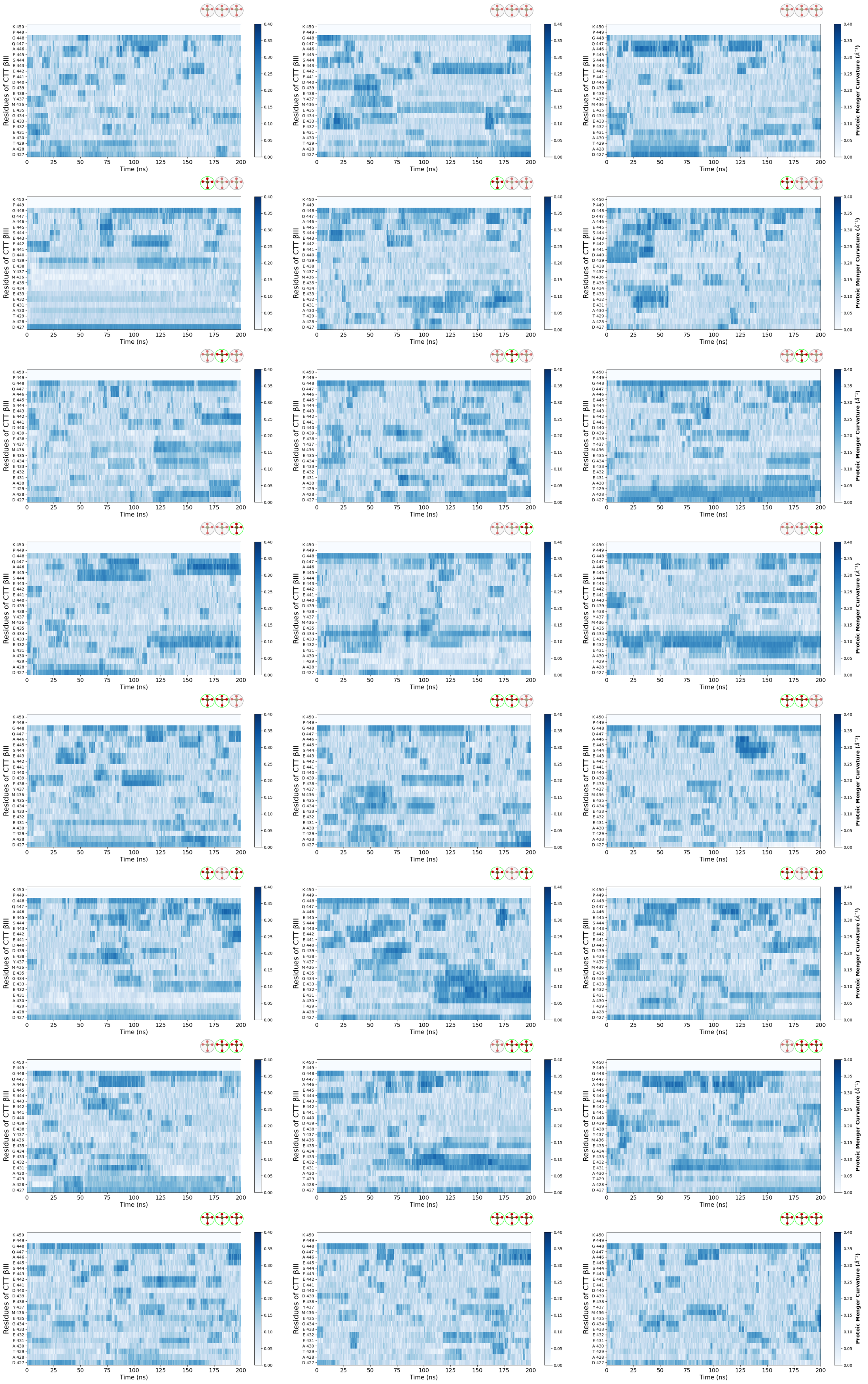

**Figure S7:** Snapshots showing the wrapping of the β-tubulin CTT (in red) around the phosphorylated tau-R2 fragment (in green). Upper panel, βI/αI/βI isotype in the P110 state. Lower panel βIII/αI/βIII isotype in the P001 state.

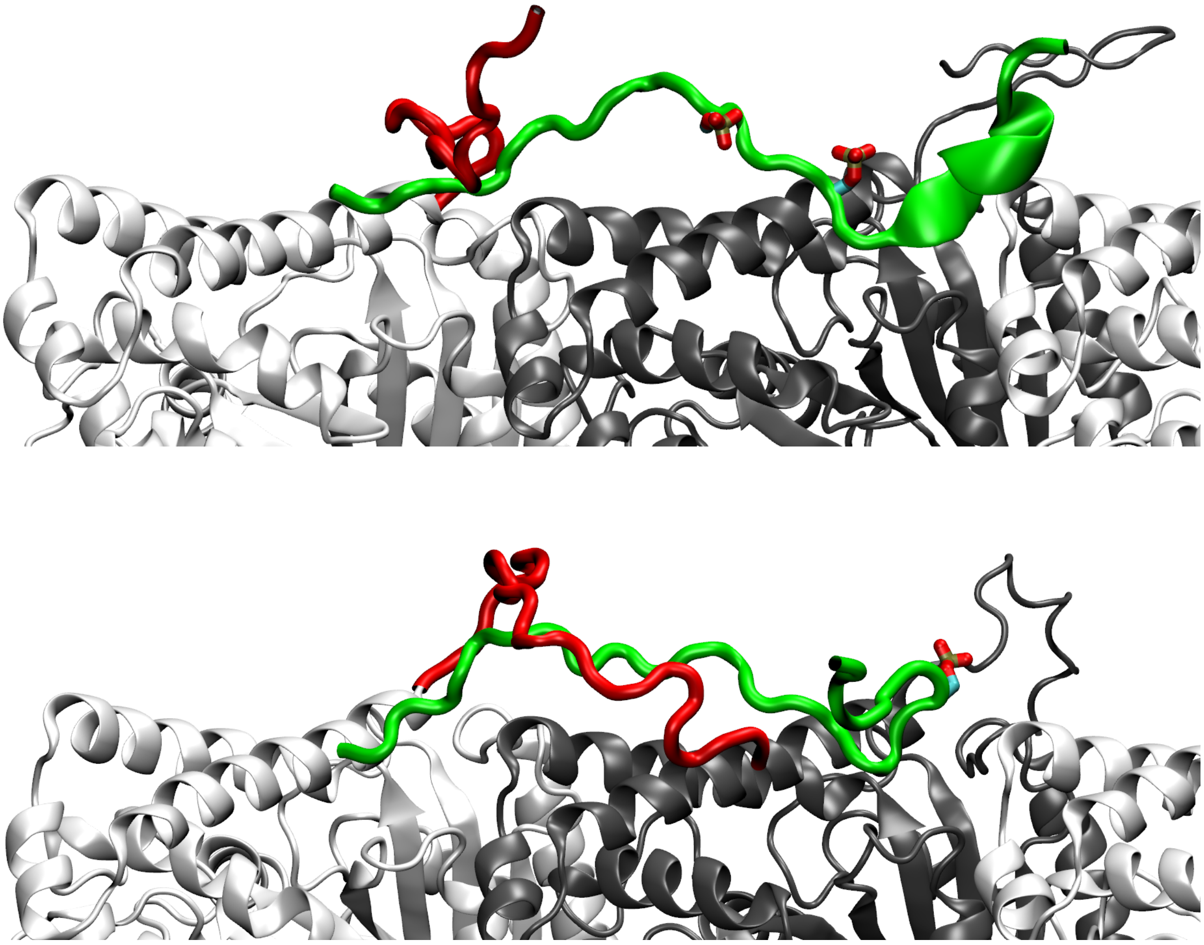
